# *In silico* screening of GMQ-like compounds reveals guanabenz and sephin1 as new allosteric modulators of acid-sensing ion channel 3

**DOI:** 10.1101/704445

**Authors:** Gerard Callejo, Luke A. Pattison, Jack C. Greenhalgh, Sampurna Chakrabarti, Evangelia Andreopoulou, James R. F. Hockley, Ewan St. John Smith, Taufiq Rahman

## Abstract

Acid-sensing ion channels (ASICs) are voltage-independent cation channels that detect decreases in extracellular pH. Dysregulation of ASICs underpins a number of pathologies. Of particular interest is ASIC3, which is recognised as a key sensor of acid-induced pain and is instrumental in the establishment of pain arising from inflammatory conditions, such as rheumatoid arthritis. Thus, the identification of new ASIC3 modulators and the mechanistic understanding of how these compounds modulate ASIC3 could be important for the development of new strategies to counteract the detrimental effects of dysregulated ASIC3 activity in inflammation. Here, we report the identification of novel ASIC3 modulators based on the ASIC3 agonist, 2-guanidine-4-methylquinazoline (GMQ). Through a GMQ-guided *in silico* screening of Food and Drug administration (FDA)-approved drugs, 5 compounds were selected and tested for their modulation of rat ASIC3 (rASIC3) using whole-cell patch-clamp electrophysiology. Of the chosen drugs, guanabenz (GBZ), an α_2_-adrenoceptor agonist, produced similar effects to GMQ on rASIC3, activating the channel at physiological pH (pH 7.4) and potentiating its response to mild acidic (pH 7) stimuli. Sephin1, a GBZ derivative that lacks α_2_-adrenoceptor activity, has been proposed to act as a selective inhibitor of a regulatory subunit of the stress-induced protein phosphatase 1 (PPP1R15A) with promising therapeutic potential for the treatment of multiple sclerosis. However, we found that like GBZ, sephin1 activates rASIC3 at pH 7.4 and potentiates its response to acidic stimulation (pH 7), i.e. sephin1 is a novel modulator of rASIC3. Furthermore, docking experiments showed that, like GMQ, GBZ and sephin1 likely interact with the nonproton ligand sensor domain of rASIC3. Overall, these data demonstrate the utility of computational analysis for identifying novel ASIC3 modulators, which can be validated with electrophysiological analysis and may lead to the development of better compounds for targeting ASIC3 in the treatment of inflammatory conditions.

## 1. Introduction

Extracellular protons modulate the activity of a wide range of ion channels and receptors, which activate sensory neurons involved in nociception and the development of pain [1]. One key group of proton sensors is the acid-sensing ion channel (ASIC) family. These voltage-independent, ligand-gated cation channels are activated by extracellular protons [2–4] and belong to the amiloride-sensitive epithelial sodium channel/degenerin (ENaC/DEG) ion channel family [5,6]. In mammals, four genes (ASIC1-4 for human and asic1-4 for rodents) encode for at least 6 different ASIC subunits (ASIC1a, ASIC1b, ASIC2a, ASIC2b, ASIC3, ASIC4), which can assemble into homo- and heterotrimeric channels displaying different pH sensitivity, current kinetics and pharmacology [7–9]. ASICs are widely expressed in the central and peripheral nervous systems [2,10] and are implicated in a range of physiological and pathological processes including nociception, mechanosensation and learning/memory [11–21]. The involvement of ASICs in such a plethora of physiological and pathological roles makes them attractive pharmacological targets for drug development and a variety of agents have been identified that act as agonists/antagonists for these channels with differing levels of selectivity [22]. Of the ASIC subunits, there is good evidence that ASIC3 is a critical acid sensor involved in acid-induced pain. ASIC3 homomers are the most sensitive to decreases in extracellular pH [7], in response to which they produce a biphasic inward current composed of a large, rapidly desensitizing transient current, followed by a smaller, non-desensitizing sustained window current (resulting from an overlap between pH-dependent activation and inactivation curves) that lasts for the duration of the acidic stimulus [23,24]. Although protons appear to be the main endogenous activators of ASICs, other molecules that modulate ASIC function have been discovered. For instance, endogenous molecules such as arachidonic acid and anandamide [25], serotonin [26], dynorphins [27] and lactate [28] all enhance ASIC3 currents in response to acidic stimulation. In addition, 2-guanidine-4-methylquinazoline (GMQ) [29], agmatine [30] and lysophosphatidylcholine [31] can activate ASIC3 at physiological pH (pH 7.4) by increasing the sustained, window current. Moreover, a range of toxins isolated from animal venoms also modulate ASIC function [32]. Among them, APETx2, a toxin isolated from the sea anemone *Anthopleura elegantissima*, inhibits the transient component of ASIC3 activation in response to an acidic stimulus (pH 4) without affecting its sustained component [33] and it has been used to establish the role of ASIC3 in a number of physiological and pathological processes, including inflammatory pain [12]. Nevertheless, the relationship between ASIC3, pain and inflammation is complex. Several histological studies, as well as those employing pharmacological ASIC3 modulation, have determined an involvement of ASIC3 in pain elicited in deep tissues such as joints, muscle and the viscera [34–40]. However, further studies using ASIC3-deficient mice have suggested a more limited role in pain [41], as well as a proposed dual role of ASIC3 in arthritis where lack of ASIC3 ameliorates pain, but increases inflammatory processes in the arthritic joint [42]. A possible explanation of these controversial results could be that inflammatory processes are sometimes [43], but not always [44], accompanied by acidosis. Nevertheless, the complex and controversial role of ASIC3 in some inflammatory processes requires the development of better pharmacological tools to dissect its precise function in such conditions. To this end, in the present study we hypothesised that we could employ GMQ as a query structure in a ligand-based *in silico* screening of FDA-approved drugs to identify novel ASIC3 modulators. From the results of the screening, we selected five different compounds with chemical and structural similarities to GMQ. From all drugs tested, (GBZ), an antihypertensive drug, caused an enhancement of acid-induced rASIC3 activation and, like GMQ, it also activated the channel at physiological pH. Given that GBZ is an agonist of α_2_-adrenoceptors [45], and with the goal of identify a more selective ASIC3 modulator, we evaluated the effect of sephin1, a GBZ derivative that has no adrenoceptor activity and may be of use in protein misfolding diseases such as multiple sclerosis [46]. Similarly to GBZ and GMQ, sephin1 activated rASIC3 at pH 7.4 and potentiated its activation in response to a mild acidosis. In summary, we demonstrate that ligand-based *in silico* approaches can be useful to identify novel small molecule modulators of ASIC3. Indeed, we have identified, from a library of FDA-approved drugs that have been proven safe for their use in humans, novel ASIC3 modulators that enhance rASIC3 activity, proving that this approach can serve to identify potential new ASICs modulating drugs that could be useful in the treatment of inflammatory disorders.

## 2. Materials and Methods

### 2.1. Ligand-based screening

The 3D structure of GMQ was obtained from pubchem (pubchem CID: 345657) and was subsequently energy-minimised using MMFF94 force field implemented in OpenBabel version 2.4.0 [47]. Using the energy-minimised GMQ structure as a 3D query, Rapid Overlay of Chemical Structures (ROCS) (version 3.2.2.2, OpenEye Scientific Software, Santa Fe, NM) [48] was used to screen a conformer library generated from eDrug3D database [49] that contains 1884 different molecular structures including structures of enantiomers and of active metabolites of FDA-approved drugs. The conformer library was generated using Omega 3.0.1.2 (OpenEye Scientific Software) [50]. For each alignment, ROCS compares 3D shape and chemical similarity and returns a Tanimoto Combo (TC) score, ranging from 0 to 2, that includes a Shape Tanimoto (maximum 1) and Colour Tanimoto (scaled colour score, maximum 1) [51]. Following manual inspection of the top 150 hits ranked by the TC score, a subset of drugs was selected for experimental testing. Molecular field-based alignment [52] of drug structures with GMQ was performed using Forge (v 10.4.2; Cresset^®^, Litlington, Cambridgeshire, UK).

### 2.2. Chinese hamster ovary cell culture and transfection

Chinese hamster ovary (CHO) cells (Sigma, passage 6 to 20) were chosen for this study due to the absence of endogenous ASIC-like currents [53] and were grown using standard procedures in the following medium: Ham’s F-12 Nutrient Mixture (Life Technologies), 10 % fetal bovine serum (Sigma), 1 % Penicillin/Streptomycin (100 U/ml, Life Technologies). 24-hours before transfecting cells, 35 mm dishes (Fisher) were coated with 100 µg/ml poly-L-lysine (Sigma) and cells from a 70-80% confluent flask were trypsinised, resuspended in 5 ml CHO medium and a volume was taken to seed cells at a 1:10 dilution (2 ml/dish). For transfections, an EGFP expression vector was used to enable identification of transfected cells and DNA was transfected at a ratio of 20:1 (rASIC3:GFP), using 1.5 µg rASIC3 DNA and 0.075 µg EGFP DNA; the transfection reagent Lipofectamine LTX (Life Technologies) was used according to the manufacturer’s protocol.

### 2.3. Whole-cell electrophysiology

Whole-cell patch clamp recordings from CHO cells were performed at room temperature 24-hours after transfection. For all the experiments, the intracellular solution contained (in mM) 110 KCl, 10 NaCl, 1 MgCl_2_, 1 EGTA, 10 HEPES, 2 Na_2_ATP, 0.5 Na_2_GTP in MilliQ water; pH was set to pH 7.3 by adding KOH and the osmolality was adjusted to 310-315 mOsm with sucrose. The extracellular solution contained (in mM) 140 NaCl, 4 KCl, 2 CaCl_2_, 1 MgCl_2_, 10 HEPES, 4 Glucose in MilliQ water; osmolality was adjusted to 300-310 mOsm with sucrose and pH was adjusted to 7.4 with NaOH. Patch pipettes were pulled from glass capillaries (Hilgenberg) using a Model P-97, Flaming/Brown puller (Sutter Instruments) and had a resistance of 4-8 MΩ. Data were acquired using an EPC10 amplifier (HEKA) and Patchmaster software (HEKA) after suitable resistance compensation. To measure the effect of the different selected compounds on rASIC3 current amplitude and inactivation time constant the following protocol was used. After 5 s of pH 7.4 solution, pH 7 or pH 6 was applied for 5 s to determine the baseline rASIC3 response. Then, in the first group of experiments, after the initial pH 7 baseline rASIC3 response, compounds were applied at pH 7 after 30 s of pH 7.4 to measure the effect of the selected compounds on pH 7 rASIC3 activation. In the second group of experiments, a second pH 6 application was performed after 10 s of pH 7.4 and 30 s of compound application to determine the effect of these compounds on rASIC3 pH 6 response. Finally, a third 5 s pH 6 or pH 7 application was performed after 30s of pH 7.4 solution to determine reversibility of any possible effect of the compounds on the channel. For concentration-response recordings of GMQ, guanabenz and sephin1 at pH 7.4 and pH 7 (sephin1), increasing concentrations of each drug (1 µM - 1 mM) were applied for 10 s with a 30 s wash period with extracellular pH 7.4 solution between each application. For pH-response recordings, extracellular solutions with a pH ranging from 7.4 to 5 with/without sephin1 were applied for 10 s with a 30 s wash period between applications. All compounds/acidic solutions were applied to cells through a gravity-driven 12-barrel perfusion system [54]. In all the experiments the holding potential was set at -60mV.

### 2.4. Molecular modelling and docking

We used three homology models of rASIC3 (Uniprot accession: O35240) based on published structures of chicken ASIC1 as the templates, solved in its closed (PDB id: 6AVE; [55]), open (PDB id: 4NTW; [56]) and desensitized state (PDB id: 2QTS; [8]) and we designated the rASIC3 models as ‘rASIC3-closed’, ‘rASIC3-open’ and ‘rASIC3-desensitized’, respectively. Detail of the model building was previously reported [57]. Selected drugs were docked to the rASIC3 models using the Lamarckian genetic algorithm (LGA) implemented in AutoDock 4.2.6 [58]. For all docking, an unbiased (“blind”) docking approach [59] was used where the entire trimers of rASIC3 models were used for generating the grid map in AutoGrid, without assuming a priori any putative binding site(s) for the ligands. The docking protocol was semi-flexible i.e. only the ligand structures were allowed to be fully flexible whilst the protein structures were held rigid. Prior to docking, structures of all drugs (obtained from PubChem) and the rASIC3 trimer were prepared using the AutoDock Tools. Five independent docking runs were performed for each drug and the pose associated with the highest reproducibility and lowest predicted free energy of interaction (ΔG, kcal/mol) was considered as the final pose for each drug. From those final docked poses of the drugs, the corresponding 2D ligand interaction diagrams were generated using PoseView™ implemented in the ProteinsPlus webserver (https://proteins.plus/). Although only the top-ranked pose for each molecule was considered for analysing the residue contacts through Poseview, representative top 1-5 poses for each drug has been shown in Fig. 7. UCSF Chimera version 1.14 (http://www.rbvi.ucsf.edu/chimera/) was used for all molecular representations.

### 2.5. Drugs

All the small molecules used in this study were purchased from Sigma (Gilliangham, UK) except for tizanidine (Tocris, Bristol, UK) and APETx2 (Smartox, Saint egrève, France). Stock solutions were made at 100 mM for tizanidine (in H_2_O), guanabenz (EtOH), cycloguanil (DMSO) and sephin1 (DMSO), 50 mM for GMQ (DMSO) and brimonidine (H_2_O), and 40 mM for guanfacine (H_2_O). For most experiments, compounds were diluted in pH 7.4 or pH 7 extracellular solution at 500 µM, however, GMQ, guanabenz and sephin1 were also diluted in extracellular pH 7.4 (or pH 7 for sephin1) solution at different concentrations. An APETx2 stock solution was made at 100 µM and diluted at 1 µM in pH 7.4 extracellular solution.

### 2.6. Data analysis

Absolute peak amplitudes were measured by subtracting the peak amplitude response from the 5-second mean baseline prior to stimulation. Peak amplitude was then normalised by dividing the absolute peak amplitude by the capacitance of the cell to obtain peak current density (pA/pF). The inactivation time constant was measured with a single exponential equation using a built-in function of Fitmaster. The sustained current to transient current ratio was calculated by measuring the size of the sustained current (peak sustained response at the end of the stimulus minus the 5- or 10-second mean baseline current; expressed as I_5s_ or I_10s_) and dividing this value by the peak amplitude current (I_5s_/I_peak_ x 100). The analysis of rASIC3 current amplitudes and kinetics was performed as previously reported [9]. Statistical analysis was performed in GraphPad Prism using a paired t-test comparing the baseline pH response (pA/pF) against the pH response after compound application for each cell. Data were plotted as a percentage of the initial pH response for each cell. In order to compare the difference between the rASIC3 activation of GMQ, GBZ and sephin1 at pH 7.4, results were expressed as a percentage of the initial pH 6 baseline ASIC3 response and unpaired t-test or one-way ANOVA with Holm-Sidak’s multiple comparison test was used. For concentration-response curves, all measurements were expressed as a percentage of the pH baseline peak current value (pH 6 or pH 7). For pH-response curves, all measurements were transformed to percent of the maximum peak current (I/Imax x 100). The EC_50_ for both concentration- and pH-response (pH_50_) experiments were determined using a standard Hill equation using GraphPad Prism. For the pH-response curve in the presence of sephin1 a biphasic equation was used in GraphPad Prism. For the analysis of pH-dependent effect of sephin1 on the rASIC3 sustained current a Gaussian distribution equation was used in GraphPad Prism. All results are expressed as mean ± standard error of the mean (SEM). All figures were made using GraphPad Prism and Adobe Illustrator CS6.

## 3. Results

### 3.1. Ligand-based *in silico* screening of novel ASIC3 modulators

Using the energy-minimised 3D structure of GMQ as a query, we used ROCS to screen a conformer library of FDA-approved drugs (eDrug3D) [49]. ROCS aligns each conformer from the target chemical library against the query or bait structure and quantifies the overall similarity between the aligned ligands as the Tanimoto Combo (TC) score. The latter is the sum of the Shape Tanimoto and the Color Tanimoto (scaled colour score), which represent the measure of similarity in 3D shape and chemical properties between the aligned moieties, respectively [51]. Of the top 150 hits ranked by the TC score, we manually inspected each individual hit for the degrees of 3D shape overlap and chemical similarity with GMQ, paying particular attention to the presence of a guanidine (or similar) moiety. This finally led us to shortlist 5 drugs, namely tizanidine (TIZ), cycloguanil (CG), brimonidine (BRI), guanfacine (GF) and GBZ (Fig. 1A). Of these hits, GBZ and GF contain an explicit (i.e. free) guanidine group (shown in red in Fig.1B), whereas CG, BRI and TIZ have an ‘implicit’ (i.e. within a ring) guanidine moiety.

**Figure 1.**
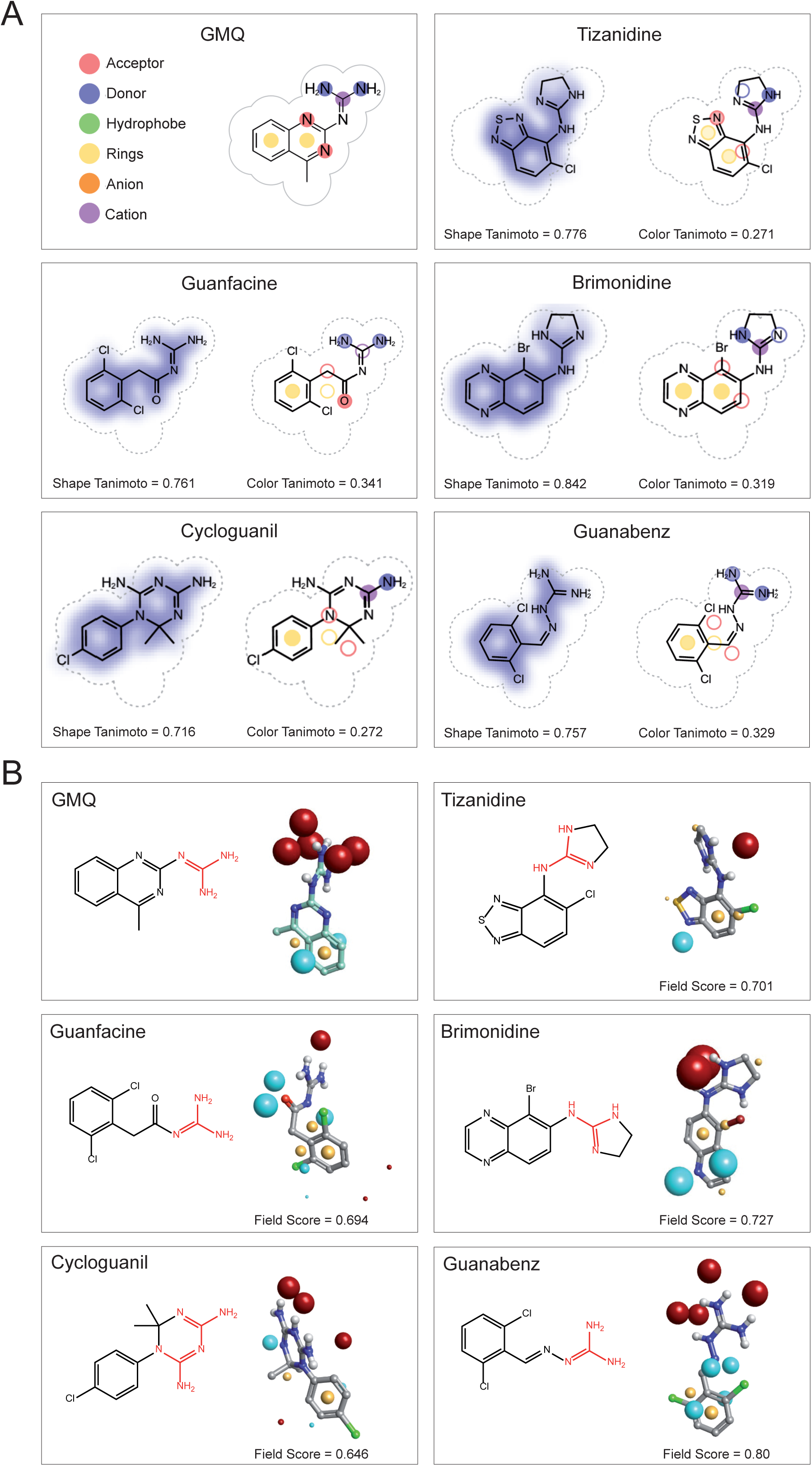
Selected FDA-approved drugs after ligand-based in silico screening using GMQ as query molecule. (A) Representations of the selected drugs showing similarity in 3D shape and chemical features with GMQ. Drugs were chosen from ligand-based in silico screening using ROCS (OpenEye Scientific Software) that ranked them on the basis of Tanimoto Combo Score. The latter is a sum of Shape Tanimoto and Colour Tanimoto score indicating similarity in 3D shape (maximum value = 1) and chemical features (maximum value = 1). The volume of the query molecule (GMQ) is shown as a dotted area and the chemical similarity aspects are shown in different colours. The figure was generated by ROCS Report (OpenEye). (B) 2D chemical structure of GMQ and selected FDA-approved drugs (Guanidine or similar moiety is shown in red) and alignment of these drugs with GMQ based on molecular electrostatic field potentials. Forge^TM^ (v 10.4.2; Cresset) was used to align the drug structures with the energy-minimised structure of GMQ. In all cases, the negative, positive and hydrophobic field points are coloured blue, red and gold, respectively (van der Waals isosurfaces are not shown). The sphere size corresponds to possible interaction strength with the cognate probe used for field point calculation. The individual molecular field similarity scores to GMQ (maximum value = 1) are given below right in each panel.

To assess how these drugs compare to GMQ in terms of overall surface electrostatics, they were aligned to the energy-minimised GMQ structure using the software Forge™ (Cresset, UK), which uses a proprietary molecular mechanics-based (’XED’) forcefield to generate and compare molecular ‘field points’ between aligned molecules (Fig. 1B). These field points represent positions of maximum interaction of a molecule with its electrostatic, steric and hydrophobic surroundings and thus effectively provides a ‘protein-centric’ view of a ligand. Upon alignment, Forge produces a field similarity score that takes both volume and molecular field points into account and a field score >0.7 is often regarded as indicator of reasonably good electrostatic similarities between the query and the bait molecule [52]. GBZ both qualitatively and quantitatively has the highest field point similarity with GMQ; in decreasing order of similarity of molecular fields, compound similarity with GMQ was computed as: GBZ > BRI > TIZ > GF > CG (Fig. 1B).

### 3.2. Effect of selected drugs on acid-induced rASIC3 activation

The following set of experiments were conducted to determine if the selected drugs modulate rASIC3 function. We performed whole-cell patch clamp recordings in CHO cells co-transfected with rASIC3 and EGFP and evaluated the effect of each one of the drugs selected on the response of rASIC3 to pH 7 and pH 6. Millimolar concentrations of GMQ are required to activate rASIC3 channels at pH 7.4, but at micromolar concentrations, GMQ sensitises pH 7-induced rASIC3 channel activation [29]. First, we evaluated the effect of each drug (500 µM) on rASIC3 when applied at pH 7. The prototypic ASIC3 agonist GMQ induced an increase in the transient (I_Peak_, Fig. 2A-B and table 1, n = 9, paired t-test, p = 0.001) sustained current (I_5s_, Fig. 2A and table 1, n = 9, paired t-test, p = 0.0012), and the ratio I_5s_/I_Peak_ (Fig 2C and Table 1, n = 9, paired t-test, p < 0.0001), suggesting a stronger effect in the sustained component of rASIC3 pH 7 activation. Among the drugs selected (summarised in Table 1 and Fig. 2A-C), TIZ and GBZ produced a similar effect, inducing and increase in all three parameters (Fig. 2A-C; TIZ: I_Peak_, n = 9, paired t-test, p = 0.0006; I_5s_, n = 9, paired t-test, p = 0.0022; I_5s_/I_Peak_ n = 9, paired t-test, p = 0.01: GBZ: I_Peak_, n = 9, paired t-test, p = 0.038; I_5s_, n = 9, paired t-test, p = 0.012; I_5s_/I_Peak_ n = 9, paired t-test, p = 0.0012). On the contrary, CG induced a slight but still significant inhibition of the transient rASIC3 current activation without affecting the sustained component or the ratio I_5s_/I_Peak_ (CG: I_Peak_, n = 8, paired t-test, p = 0.0061; I_5s_, n = 8, paired t-test, p = 0.2; I_5s_/I_Peak_ n = 8, paired t-test, p = 0.15). GF and BRI induced contrary effects on the I_5s_/I_Peak_ ratio, whilst GF produce an enhancement of this parameter, suggesting a stronger enhancement of the sustained component, BRI induced an inhibition (GF: I_5s_/I_Peak_ n = 9, paired t-test, p = 0.03; BRI: I_5s_/I_Peak_ n = 6, paired t-test, p = 0.007).

**Figure 2.**
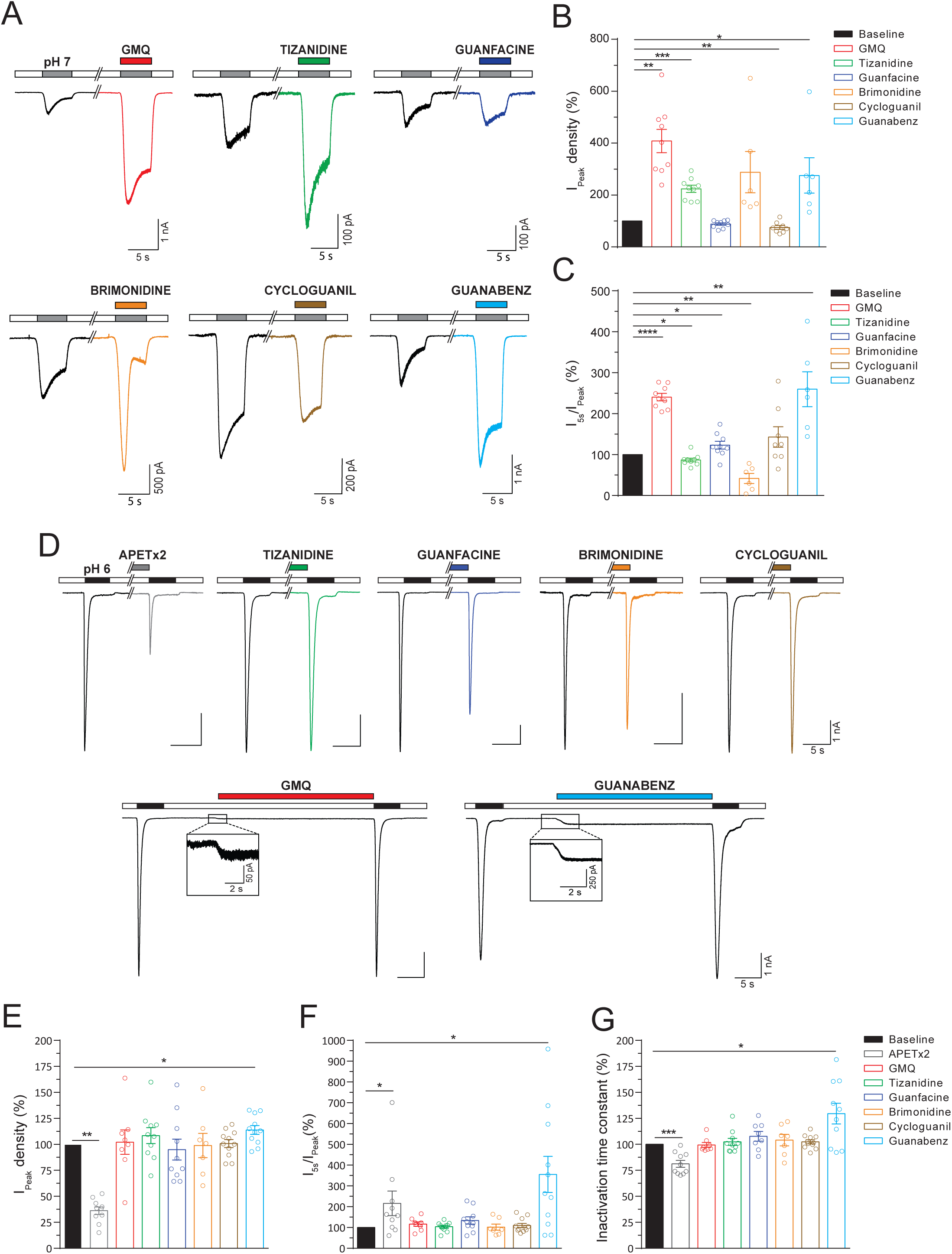
Effect of the selected drugs rASIC3 response to pH 6. (A) Example traces of the effect for the 5 different selected drugs (500 µM) on pH 7-induced rASIC3 activation together with example trace showing the effect of the prototypic rASIC3 modulator GMQ (500 µM). In all cases drugs were applied at pH 7. (B-C) Bar plots showing the effect of the selected drugs and GMQ on transient (peak) (B) and the sustained current (expressed as the ratio I_5s_/I_peak_) (C). (D) Example traces of the effect for the 5 different selected drugs (500 µM) on pH 6-induced rASIC3 activation together with example traces showing the effect of known rASIC3 modulators, APETx2 (1 µM) and GMQ (500 µM). In all cases drugs were applied for 30 s prior to pH 6-induced rASIC3 activation. (B-D) Bar plots showing the effect of the selected drugs, APETx2 and GMQ on transient (peak) (B) and sustained current (C), and inactivation time constant (D) of rASIC3 activation. All values were normalised to baseline pH 7 or pH 6 rASIC3 activation and expressed as means ± SEM (n = 6-11, paired t-test, *p <0.05 and ***p ≤ 0.005 vs baseline pH 7 or pH 6 activation).

**Table 1.**
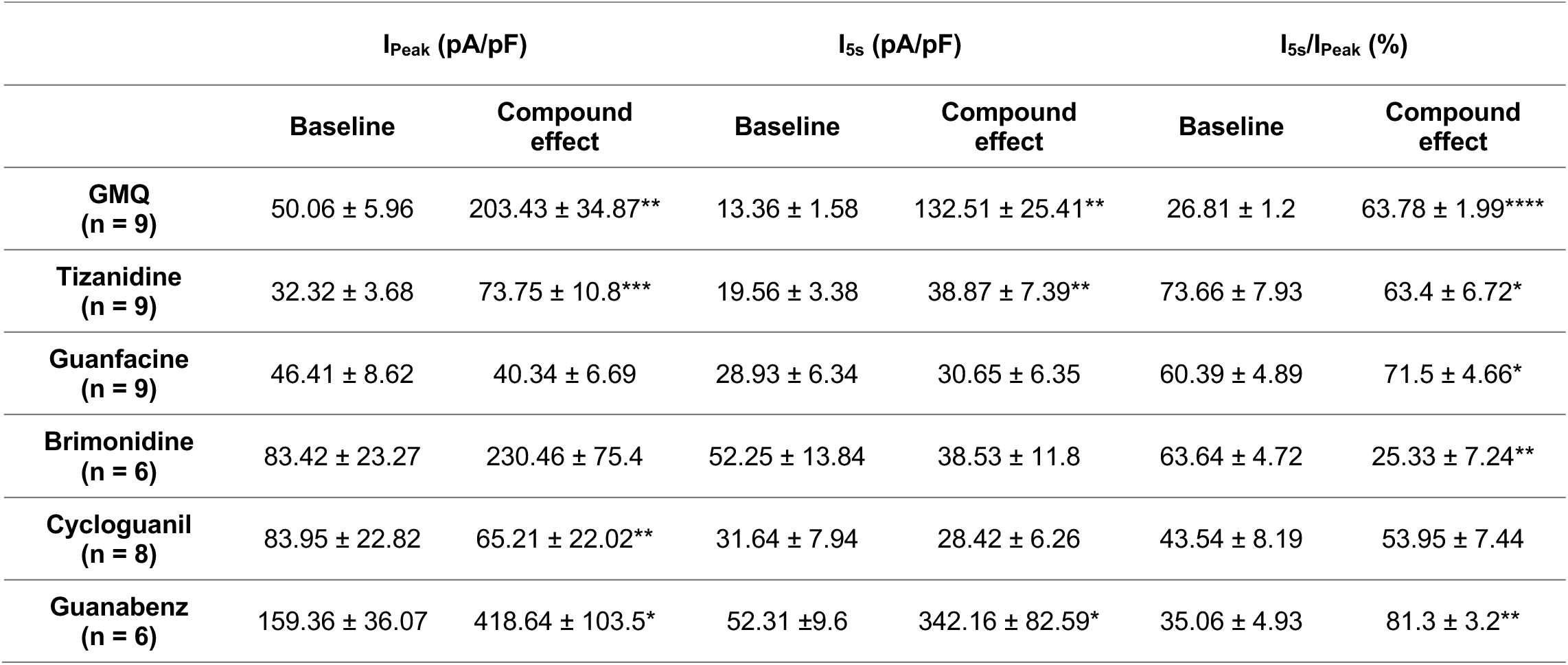
Effect of the selected compounds applied at pH 7 on rASIC3 activation. Comparison of the mean ± SEM of the amplitude of the transient (I_peak_) and sustained current (I_5s_), together with the ratio I_5s_/I_peak_ between baseline rASIC3 pH 7 response and after application of each compound at pH 7 (Paired t-test, *p ≤ 0.05, **p ≤ 0.01, ***p ≤ 0.005, ****p ≤ 0.0001).

We next tested the effect of the pre-application (30 s) of each drug (500 µM) on the rASIC3 response to pH 6. In this series of experiments, we also evaluated the effect of GMQ and APETx2, which were used as positive controls. As expected, in rASIC3-expressing CHO cells, APETx2 (1 µM) application did not activate the channel at physiological pH (pH 7.4), but produced a significant inhibition of transient (I_Peak_, Fig. 2D-E, n = 10, paired t-test, p = 0.0013) and sustained (I_5s_ in Fig. 2A, n = 10, paired t-test, p = 0.029) current evoked by pH 6 (Table 2). However, the ratio I_5s_/I_Peak_ was significantly increased (Fig 2F and Table 2, n = 10, paired t-test, p = 0.036), indicating that the predominant APETx2 inhibitory effect is exerted on the transient phase as previously described [33]. In addition, APETx2 significantly accelerated the inactivation time constant (Tau) of rASIC3 (Fig. 2G and Table 2, n = 10, paired t-test, p = 0.0005). By contrast, GMQ generated a sustained inward current at pH 7.4 as reported previously [29], but did not significantly modulate channel current amplitude or inactivation kinetics (Table 2).

**Table 2.**
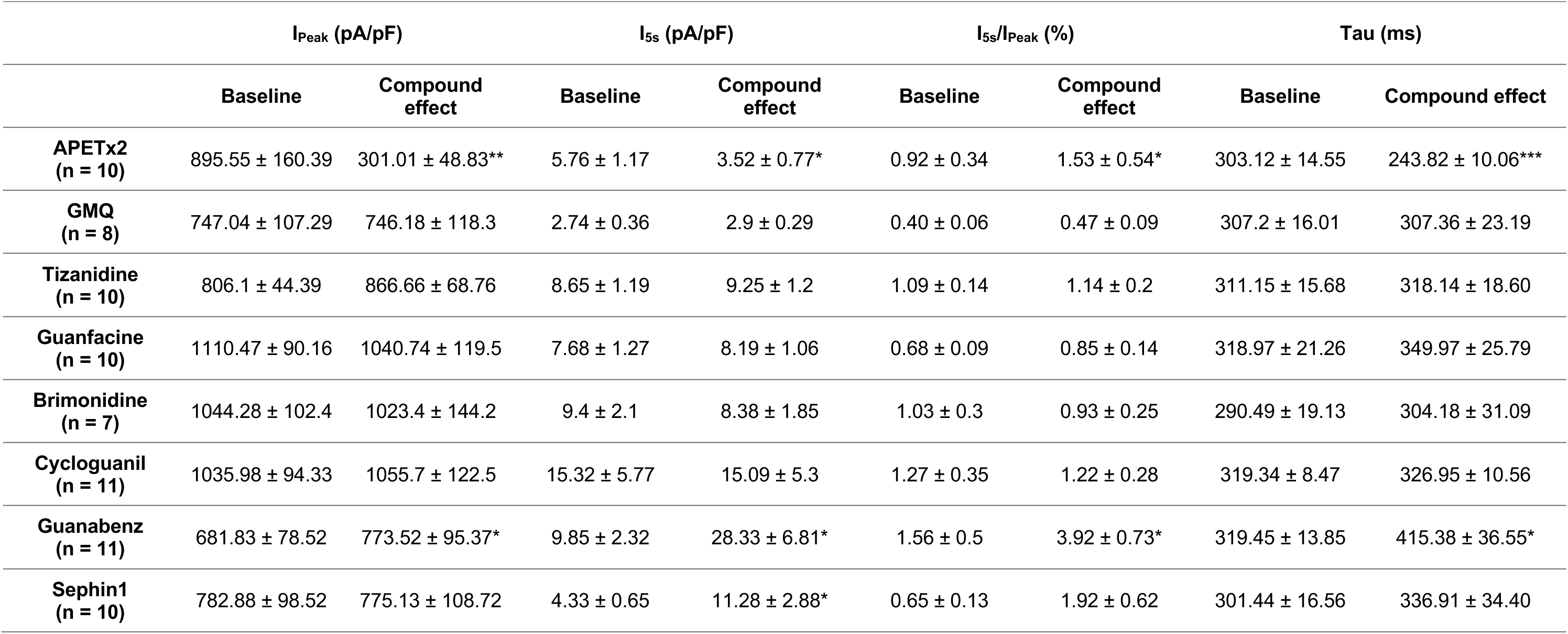
Effect of the preapplication of the selected compounds on rASIC3 current kinetics. Comparison of the mean ± SEM of the amplitude of the transient (I_peak_) and sustained (I_5s_) current, together with the ratio I_5s_/I_peak_ and the inactivation time constant (Tau) between baseline rASIC3 pH 6 responses and after compound application (Paired t-test, *p ≤ 0.05, **p ≤ 0.01, ***p ≤ 0.005).

Among the 5 drugs tested in these series of experiments (summarised in Table 2), with the exception of GBZ, none of them produced a significant change on rASIC3 current amplitude or inactivation kinetics (Fig. 2E-G). Unlike all other compounds tested, and in a similar fashion to GMQ, we observed that GBZ, a drug currently used to treat hypertension, activated rASIC3 at physiological pH (Fig. 2D). However, unlike GMQ, pre-application of GBZ elicited a significant increase in the transient and sustained components of the pH 6-induced rASIC3 current (Fig. 2E, I_Peak_, n = 11, paired t-test, p = 0.0173; I_5s_, n = 11, paired t-test, p = 0.015). The stronger potentiating effect upon the sustained current (303%) compared with the effect on the transient current (13%) produced a significant increase in the I_5s_/I_Peak_ ratio (Fig. 2F, n = 11, paired t-test, p = 0.016) and the inactivation time constant of the pH 6-induced rASIC3 response was significantly increased by GBZ (Fig. 2G, n = 11, paired t-test, p = 0.014).

### 3.3. GBZ activates rASIC3 at physiological pH and likely binds to the nonproton ligand sensor domain

Given that GBZ was the only selected drug able to activate rASIC3 at pH 7.4 and potentiate its response to pH 7 and pH 6, we decided to investigate further the mechanism action of GBZ on rASIC3. In order to quantify the rASIC3 activation at physiological pH observed by GBZ, we performed electrophysiological recordings and compared the results to the effects observed with GMQ. At pH 7.4, even though both GMQ and GBZ (1 mM), are capable of activating rASIC3, the activation of rASIC3 by GBZ is of significantly of smaller magnitude compared to GMQ-induced rASIC3 activation (10.8 ± 1.76 % for GMQ vs 5.06 ± 1.04 for GBZ, Fig.3A and 3B, unpaired t-test, n = 12-13, p = 0.0061).

**Figure 3.**
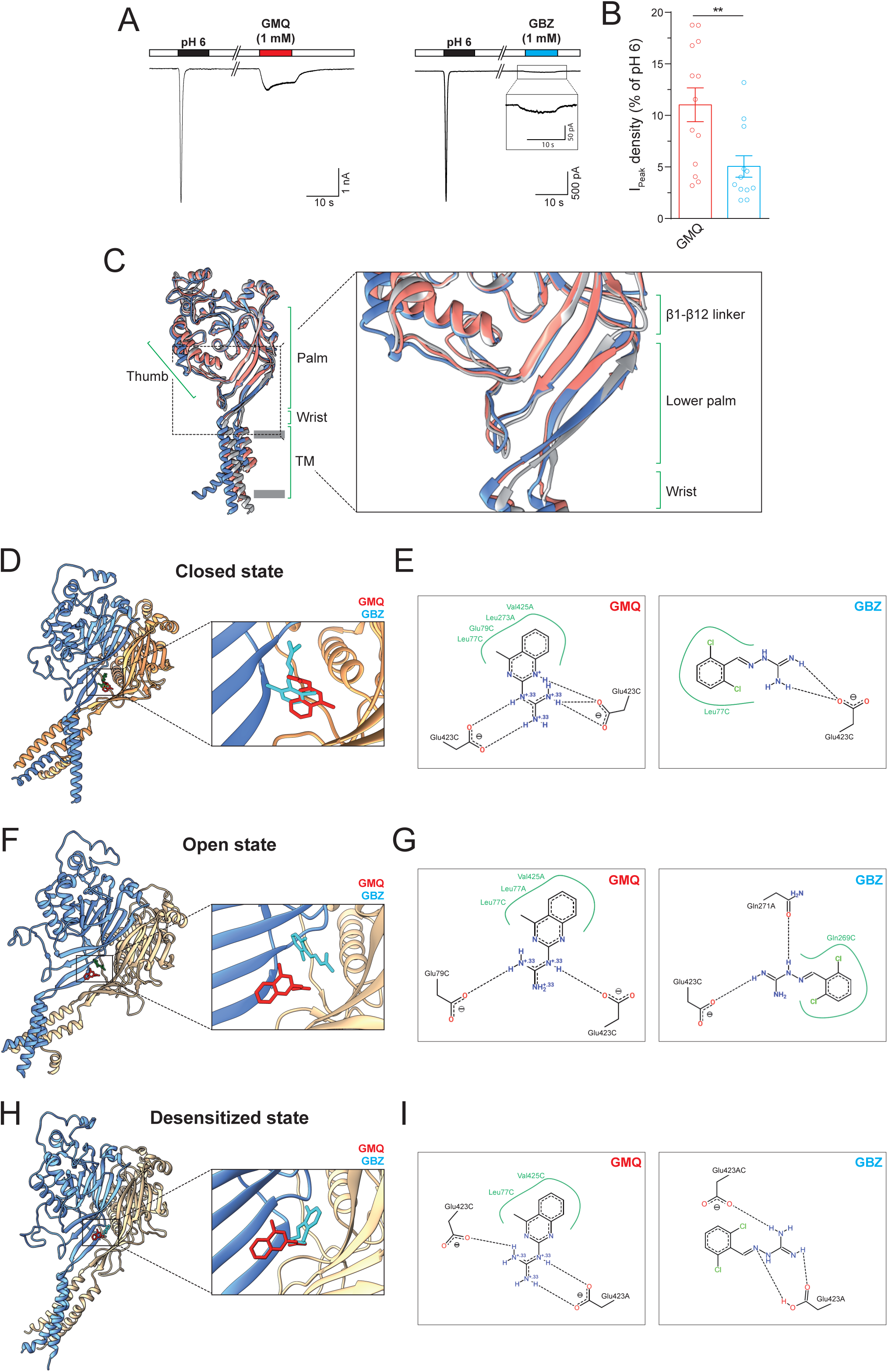
Effect of GBZ on rASIC3 response at physiological pH and proposed binding site and binding mode of GBZ on rASIC3 closed, open and desensitized models. (A) Example traces of rASIC3 activation at physiological pH (pH 7.4) elicited by GMQ (1 mM) and GBZ (1 mM). Inset showing magnification of GBZ activation. (B) Bar plot showing the quantification of GMQ- and GBZ-induced rASIC3 activation at pH 7.4 normalised against ASIC3 pH 6 response (n = 12-13, unpaired t-test, **p ≤ 0.01). (C) The structures of chicken ASIC1a representing the closed (PDB id: 6AVE, coloured in pale blue), open (PDB id: 4NTX, coloured in salmon) and desensitized (PDB id: 4NYK, coloured in grey) are shown in cartoon representations and were overlaid using UCSF Chimera 1.14. Various regions are indicated in canonical terms. (D, F and H) Representative top-ranked poses of GMQ (red) and GBZ (green) obtained through AutoDock 4.2.6-based blind docking against rASIC3-closed, rASIC3- open and rASIC3-desensitized models, respectively, are shown. For clarity, only two subunits of the channel are shown in cartoon representations (subunit A in golden colour and subunit C in pale blue colour). A close-up view of the site and the docked poses for each model is shown to the left. (E, G and I) PoseView™ derived ligand interaction diagrams for the docked poses of GMQ and GBZ are shown. A and C indicate the two subunits of rASIC3.

Given the structural and chemical similarities between GMQ and GBZ (Fig. 1), we next performed docking experiments to elucidate the binding mechanism of GBZ to rASIC3. The carboxyl-carboxylate interaction pair formed by the residues E79 and E423 in the palm domain of rASIC3 has been implicated in GMQ binding and the cavity where these two amino acids are localised has been named the ‘nonproton ligand sensor domain’ because several ‘nonproton’ ASIC3 ligands such as GMQ, agmatine and serotonin all bind this domain [26,29,30]. Although GBZ shares certain structural and chemical properties with GMQ, including a guanidine moiety (Fig. 1B), these molecules may or may not share the same binding site or manifest similar binding mode to the same site on ASIC3. Besides, the precise binding mode of any ‘nonproton’ ASIC3 ligand including GMQ to the nonproton ligand sensor domain is likely to be influenced by the state of ASIC3, since some local structural changes, though subtle, can be observed in the lower palm region of overlaid chicken ASIC1 structures solved in closed, open and desensitized states (Fig. 3C) and this has been highlighted in a recent review [60].

To address all these aspects, we performed *in silico* blind docking experiments with GMQ and GBZ against our three rASIC3 homology models (designated as rASIC3-closed, rASIC3-open and rASIC3-desensitized) that were made using template chicken ASIC1 structures solved at closed [55], open [56] and desensitized states [8], respectively. In our unbiased docking approach, GMQ preferentially docked to a pocket located in the palm domain of all three rASIC3 models (Fig.3D, 3F and 3H) which has been computationally and experimentally established in previous studies as the likely binding site for GMQ and designated as the nonproton ligand sensor domain [29,61]. However, subtle differences in its proposed binding mode were noticed in rASIC3-closed (Fig. 3D), rASIC3-open (Fig. 3F) and rASIC3-desensitized (Fig.3H) models. The 2D ligand interaction diagrams for GMQ docked to those rASIC3 models show 4-methylquinazoline moiety makes a hydrophobic interaction with L77 and V425 in all 3 rASIC3 models but for rASIC3-closed model (Fig 3E), two additional residues namely E79 and L273 seem also to participate. Interestingly, the guanidinium moiety of GMQ seems to form two salt bridges with two E423 residues present in the same or in different monomeric subunits in case with rASIC3-closed (Fig. 3E) and rASIC3-desensitized (Fig. 3I) structures, respectively whilst for the rASIC3-open model, E423 of one subunit is replaced by E79 in making such salt bridge (Fig. 3G). GBZ also docked to the same location and it broadly retains similar interactions (Fig. 3E, 3G and 3I). Like GMQ, its mode of interaction also seems to vary depending on the rASIC3 model. For example, its guanidinium moiety makes two salt bridges with E423 of a single monomer (for rASIC3-closed, Fig. 3E), E423 and Q271 of different monomers (for rASIC3-open, Fig. 3G) and E423 from two different monomers (for rASIC3-desensitized, Fig. 3I). Also, no or fewer hydrophobic contacts were noticeable for GBZ in all rASIC3 models (L77 in rASIC3-closed model and Q269 in rASIC3-desensitized model), presumably due to lack of an additional aromatic ring when compared to GMQ structure (Fig. 3E, 3G and 3I).

Taken together, these results indicate that GBZ modulates rASIC3 in the mild acidic range and that its binding site likely overlaps with that of GMQ, the so called nonproton ligand sensor domain. However, their precise modes of binding may be different and can be subjected to further variations depending on the state of the channel, which may underlie the different potency and efficacy observed on rASIC3 current activation.

### 3.4. The GBZ derivative, sephin1, positively modulates rASIC3

Recently, it has been demonstrated that a GBZ derivative without α_2_-adrenoceptor activity, sephin1 (also known as IFB-088; Fig.4A) acts as an inhibitor of a regulatory subunit of the stress-induced protein phosphatase 1 (PPP1R15A) [62]. Sephin1 is effectively the mono-chlorinated version of GBZ and is currently under investigation in clinical trials and hence was not included in the eDrug3D database that we initially screened with ROCS using GMQ as bait. ROCS-based alignment of sephin1 with GMQ revealed by far the highest similarity among the 5 initially selected FDA-approved compounds in terms of both 3D shape (>90%) and chemical features (>50%) with an overall TC score of 1.481 (Fig. 4A). In agreement with this, sephin1 appeared to be the most similar to GMQ among all FDA-approved selected drugs in terms of the molecular field points with the highest overall field score (Field Score = 0.847, Fig. 4B).

**Figure 4.**
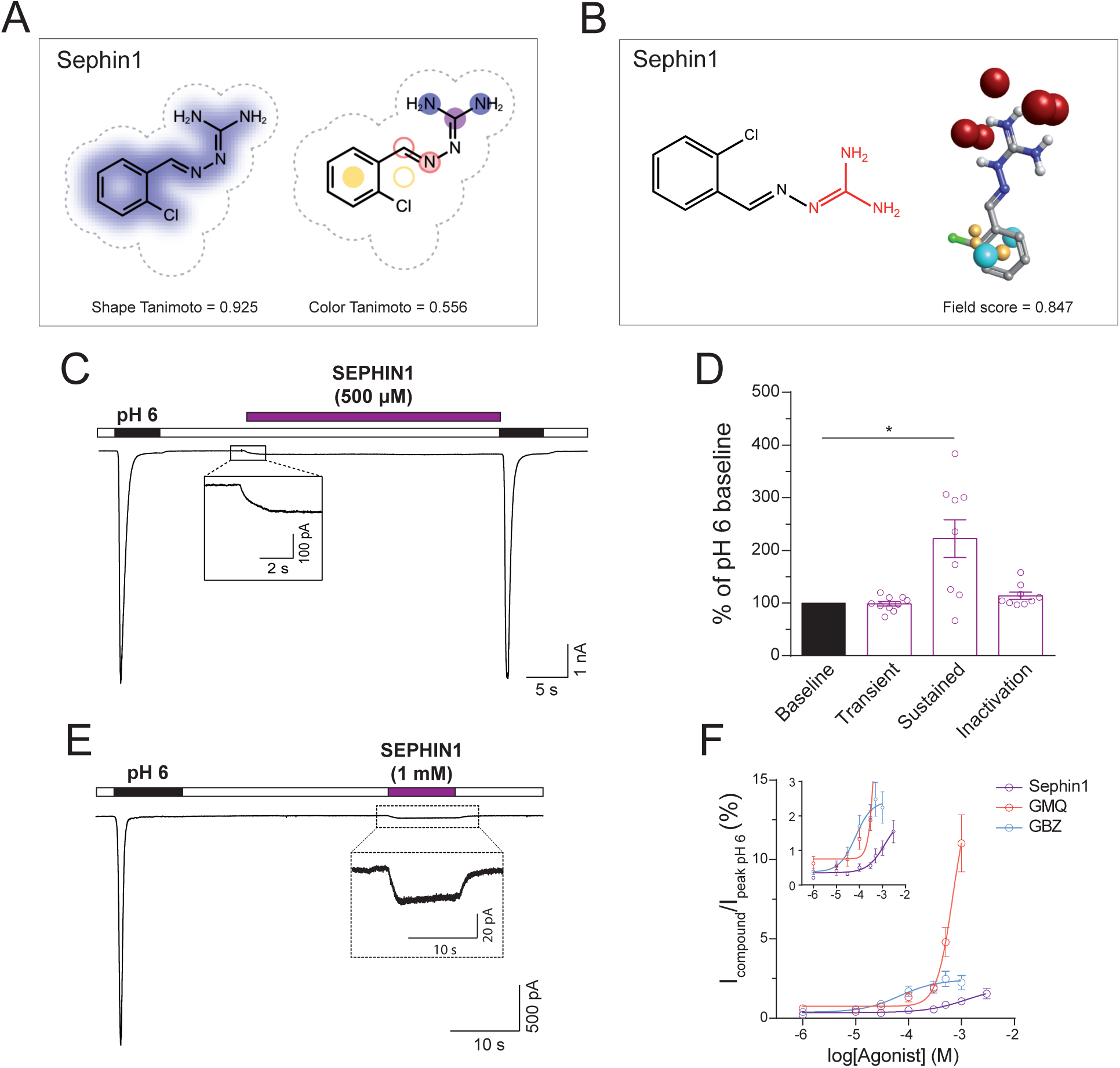
Effect of sephin1 on rASIC3 activation. (A-B) Representation of sephin1 showing similarity in 3D shape and chemical features with GMQ (A) and molecular field electrostatics (B) as described in Fig.1 for selected drugs. (C) Example trace of effect of sephin1 (500 µM) on pH6-induced rASIC3 activation. (C inset) Magnification of rASIC3 activation induced by sephin1 at pH 7.4. (C) Bar plot quantifying sephin1 effect on the transient and sustained normalised current and inactivation time constant of the pH 6-induced rASIC3 activation expressed as a percentage of the pH 6 baseline activation (n = 10, paired t-test; *p <0.05 vs baseline pH 6 activation). (E) Example trace showing rASIC3 activation by sephin1 at pH 7.4. (E inset) Magnification of the current elicited by sephin1 (n = 12). (F) Concentration-response effect of increasing concentrations of GMQ, GBZ and sephin1 (1 µM–1 mM) applied at pH 7.4 on rASIC3. Absolute values were normalised using the capacitance of each cell (pA/pF) and expressed as a percentage of the pH 6-induced rASIC3 activation. A non-linear regression using a sigmoidal function was used to determine the EC_50_ of all three molecules on rASIC3 (n = 4-12, GMQ EC_50_ = 0.68 mM; GBZ EC_50_ = 67.5 µM; sephin1 EC_50_ = 1.22 mM). (F inset) Magnification of the concentration-response curves of all three drugs.

Given the high molecular similarity observed *in silico* for sephin1 and GMQ/GBZ, we evaluated the effect of sephin1 on the acid-induced rASIC3 response. Firstly, we observed that sephin1 (500 µM) did not affect the transient component of pH 6-induced rASIC3 activation, similar to the effect observed for GMQ but unlike GBZ (Fig. 2E and table 2), however it did induce an increase in the amplitude of the sustained component (I_peak_, Fig. 4C-D and table 2, n = 10, paired t-test, p = 0.82; I_5s_, Fig. 4C-D and table 2, n = 10, paired t-test, p = 0.03) without significantly affecting the I_5s_/I_peak_ ratio (I_5s_/I_peak_, Fig. 4C-D and table 2, n = 10, paired t-test, p = 0.13) or the inactivation time constant (Tau, Fig. 4C-D and table 2, n = 10, paired t-test, n = 10, p = 0.09, Fig. 4C). When comparing the effect of sephin1, GBZ and GMQ on the pH 6 ASIC3 response, we did not observe significant differences in the transient phase of ASIC3 activation, however, we did observe a significantly bigger increase in the sustained component after GBZ in comparison to GMQ (GMQ, 11.4 ± 7.96 % vs GBZ, 303.07 ± 107.1 %, F(2,26) = 3.420, t(26) = 2.567, ANOVA, Holm-Sidak’s multiple comparison test, p = 0.0483). Similarly to GMQ and GBZ, 500 µM and 1 mM sephin1 also activated rASIC3 at pH 7.4 (Fig. 4C and 4E insets), however, sephin1’s activation, as with that observed for GBZ, was significantly smaller than GMQ ASIC3 activation at pH 7.4 (GMQ, 10.8 ± 1.76 % vs GBZ, 5.06 ± 1.04 %, F(2,34) = 5.673, t(34) = 3.423, ANOVA, Holm-Sidak’s multiple comparison test, p = 0.0033; GMQ, 10.8 ± 1.76 % vs sephin1, 3.32 ± 0.8 %, F(2,34) = 5.673, t(34) = 4.42, ANOVA, Holm-Sidak’s multiple comparison test, p = 0.0003). In order to compare the efficacy and potency of all three molecules on their action on rASIC3 at pH 7.4, we performed a concentration-response (CR) analysis. Due to a limitation in the solubility of the drugs in their solvent, we could not perform a full CR experiment, but nevertheless, we could compare the effect of all three molecules estimating the lowest EC_50_ value. GMQ appears to be the most efficacious of all three molecules, showing an activation of 10.8 ± 1.76 % at 1 mM (% of pH 6 initial response) (Fig. 4F), whereas at the same concentration of GBZ and sephin1 elicited an activation of 5.06 ± 1.04 % and 3.32 ± 0.8 %, respectively (Fig. 4F inset). On the other hand, GBZ appears to be the most potent of all three molecules with an estimated EC_50_ of 67.5 µM, compared to 0.68 mM and 1.22 mM for GMQ and sephin1, respectively (Fig. 4F inset). Even though sephin1 appears to be the least potent and efficacious of all three molecules, like GMQ and GBZ, sephin1 induced a strong sensitisation, in a concentration-dependent fashion, of the transient and sustained (I_10s_ in Fig.5D and 5H) components of the pH 7-induced rASIC3 activation (Fig. 5A-D), revealing an EC_50_ of 28.93 µM for the rASIC3 transient current at pH 7 (Fig. 5B). At 500 µM, like GMQ and GBZ, sephin1 also enhanced the transient and sustained currents of rASIC3 (I_peak_ 87.58 ± 37.33 pA/pF vs I_peak sephin1_ 230.84 ± 80.42 pA/pF n = 9, paired t-test, p = 0.016; I_5s_ 27.13 ± 18.65 pA/pF vs I_5s sephin1_ 148.37 ± 64.98 pA/pF, n = 9, paired t-test, p = 0.03; I_5s_/I_peak_ 30.24 ± 6.8 % vs I_5s sephin1_ 65.19 ± 6.54 % n = 9, paired t-test, p < 0.0001), however, we found no significant differences in the sensitisation of ASIC3 pH 7 response between all three molecules. Given the high similarities in the action of sephin1 and GMQ, we hypothesized that sephin1 also binds to the nonproton ligand sensor domain of rASIC3. We next tested the pH dependency of rASIC3 modulation by sephin1. The application of different pH solutions ranging from pH 7.4 to 5 (Fig. 5G and 5H) on rASIC3 induced currents of increasing magnitude to produce a sigmoidal curve that could be fitted to reveal a pH_50_ value of 6.36 (Fig. 5G inset), in accordance with previous reports [7,9]. However, in the presence of sephin1 (500 µM), rASIC3 transient activation followed a pH-dependent, biphasic curve (Fig. 5H inset), suggesting an interplay between the action of sephin1 on rASIC3 and its proton activation. As showed previously by Yu et al. using GMQ [29], sephin1 activated rASIC3 at pH 7.4, but also potentiated its sustained current at all pH solutions tested, showing a Gaussian distribution with a peak at pH 6.56 (r^2^ = 0.43, Fig. 5I). Similarly to GBZ, *in silico* blind docking experiments against all rASIC3 models suggest that the E423 but not the E79 of the nonproton ligand sensor domain engages in salt bridge interaction with the guanidinium moiety of sephin1. The aromatic ring of sephin1 can have possible hydrophobic interactions with residue A378, L77, E423 and E269 (Fig. 6B, 6D and 6F), depending on the nature of the rASIC3 model. Altogether, these results show that sephin1 modulates rASIC3 activation possibly through interacting with the nonproton ligand sensor domain of the channel.

**Fig. 5.**
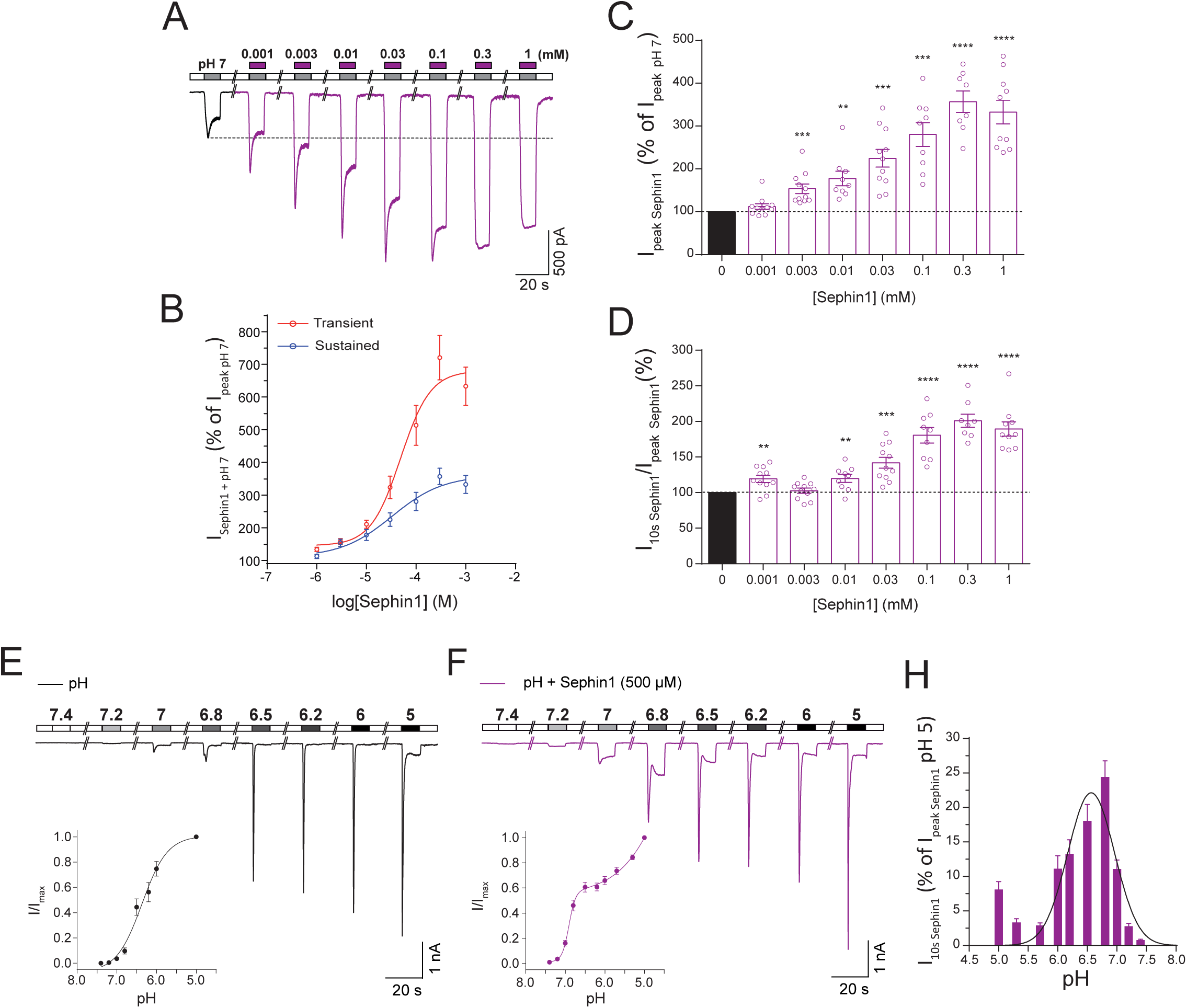
Effect of sephin1 on rASIC3 response to mild acidosis (pH 7) and pH-dependency. (A) Concentration-response effect of increasing concentrations of sephin1 (1µM–1mM) applied at pH 7 on rASIC3. (B) Fitting of normalised values for the transient (peak) and sustained current (pA/pF) obtained in A using a sigmoidal function. (C and D) Bar plot showing the quantification of the data points obtained in A for each sephin1 concentration. Normalised values were expressed as a percentage of the baseline rASIC3 pH 7 activation (n = 8-11, paired t-test, **p ≤ 0.01, ***p ≤ 0.005, ****p ≤ 0.001, vs baseline pH 7 activation). (E and F) pH-dependent effect of sephin1 on rASIC3. Example traces of rASIC3 response to a pH range from 7.4 to 5 with and without sephin1 (500 µM) (n = 7-11) (E and F insets) pH-response curves of rASIC3 activation by different pH extracellular solutions used with (E inset) and without (F inset) sephin1 (500 µM). (H) Bar plot showing the quantification of the normalised sustained current (pA/pF) elicited by sephin1 at different pH solutions.

**Fig. 6.**
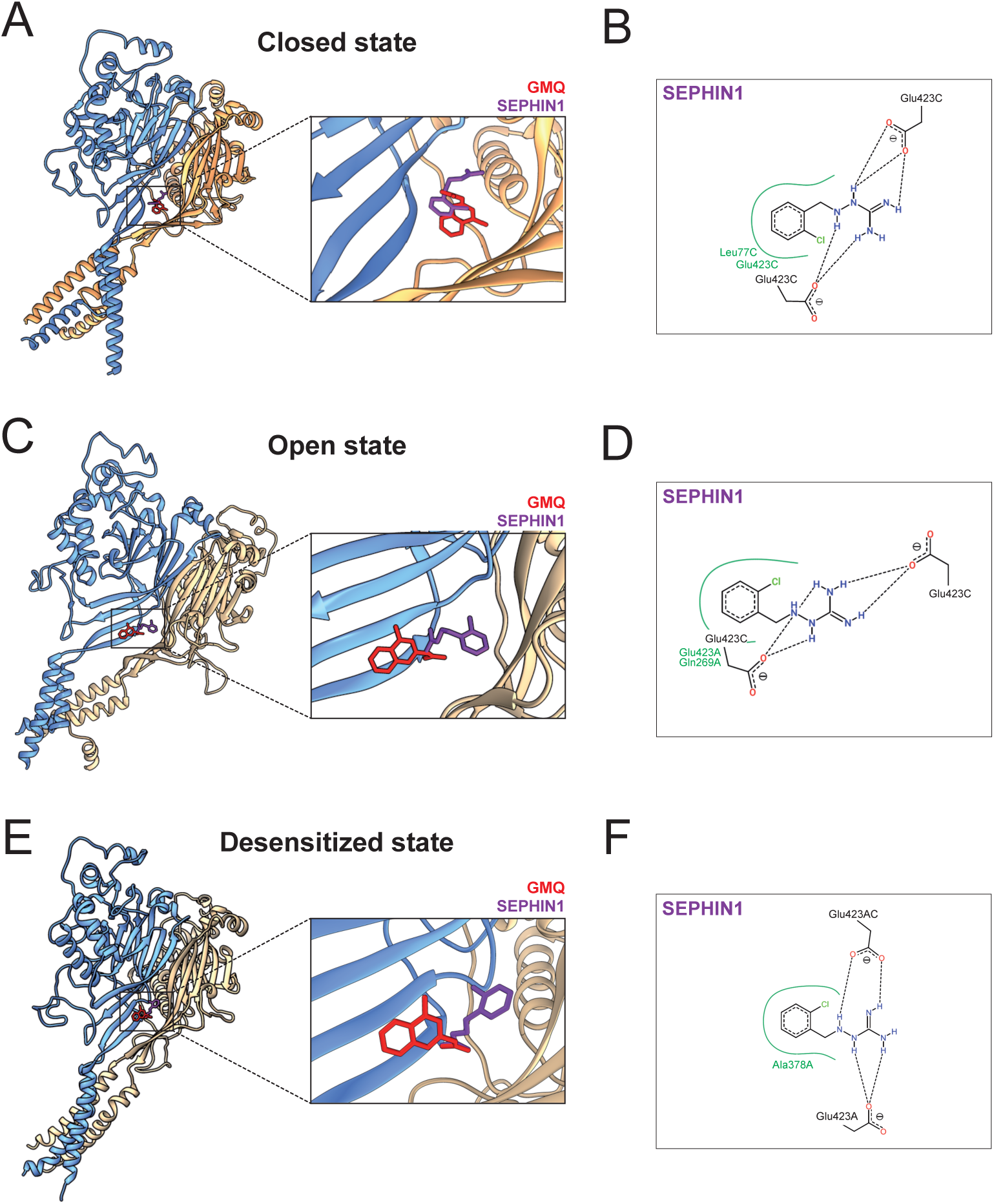
Proposed binding site and binding modes of Sephin1 on rASIC3 models representing possible closed, open and desensitized conformations. (A, C and E) Representative top-ranked poses of GMQ (red) and sephin1 (purple) obtained through AutoDock 4.2.6-based blind docking against rASIC3-closed, rASIC3-open and rASIC3-desensitized models are shown. For clarity, only two subunits of the channel are shown in cartoon representations (subunit A in golden colour and subunit C in pale blue colour). A close-up view of the site and the docked poses is shown to the left. (B, D and F). PoseView™-derived ligand interaction diagrams for the docked poses of sephin1 and GMQ for each model are shown. A and C indicate the two subunits of rASIC3-closed model.

## 4. Discussion

The ASIC family of ion channels have been implicated in many physiological and pathological processes including nociception [14,19,63], where they have been established as attractive pharmacological targets for treating pain. ASIC3 is of particular interest given its high expression in primary sensory neurones [10] and its involvement in inflammatory pain originating from different tissues including muscle, joints and skin [12,34,35,64–66]. Therefore, the exploration of novel ASIC3 modulators could increase our knowledge of ion channel function and also be pivotal for the development of new strategies to counteract the detrimental effects of dysregulated ASIC3 activity in pathophysiological states. Many molecules that modulate ASIC3 function have been discovered, ranging from non-selective ASIC3 blockers such as amiloride that acts as a pore blocker and paradoxically stimulates ASIC3 at pH 7.4 [3], to more specific molecules, such as the inhibitory toxin APETx2, which inhibits the acid-induced transient ASIC3 current [33] but also inhibits Na_V_1.8 channels [67], and the agonist GMQ, which activates ASIC3 at pH 7.4 and potentiates its activation in response to an acidic stimulus through the nonproton ligand sensor domain of ASIC3 [29,61]. However, GMQ is not a specific agonist of ASIC3 as several studies have demonstrated that GMQ also modulates other ASIC subunits [68,69] as well as GABA_A_ receptors [70]. In the present study, we used a ligand-based *in silico* screening of FDA-approved drugs to identify novel rASIC3 modulators. Of the top 150 hits ranked by TC score, we selected 5 different drugs with the highest structural and chemical resemblance to GMQ (Fig. 1A), including the presence of a guanidine group. Using an independent algorithm implemented in Cresset’s Forge™, we then aligned these 5 drugs with GMQ and compared their surface electrostatic properties represented by the molecular field points. Of the selected drugs, GBZ showed a striking resemblance to GMQ with regard to surface electrostatics and this was reflected in the highest observed value for the field score (0.80). However, the surface electrostatic analysis of sephin1 resulted in an even higher resemblance to GMQ with a field score of 0.847. Of the remaining 4 drugs, only BRI, which had the 2^nd^ best field score, showed some degree of electrostatic similarity with GMQ (Fig.1B).

We next sought to experimentally evaluate the effects of these drugs on acid-induced rASIC3 activation. Several drugs showed a modulation of the rASIC3 pH 7 response (Fig. 2A-C), however, among them only one showed a modulatory effect on rASIC3 acid response to pH 6, namely GBZ, an α_2_-adrenoceptor agonist. When compared to GMQ, GBZ exhibited a high TC score (1.086) (Fig. 1B) and the highest molecular field score (0.8) among the FDA-approved drugs selected (Fig. 1), together with the presence of an explicit guanidine group. Interestingly, TIZ that showed a higher TC score (Fig. 1A; 1.047) and a lower field score (Fig. 1B) in terms of surface electrostatics than GBZ did enhance only the pH 7 but not the pH 6 rASIC3 response. The precise reason for such behaviour of TIZ remains to be properly investigated but it indicates that a free guanidinium moiety is perhaps important to influence the pH 6 ASIC3 response which TIZ clearly lacks, unlike GBZ and GMQ. Indeed, a pseudo or implicit form of the guanidinium moiety contained in the 4,5-dihydro-1*H*-imidazol-2-amine ring of TIZ appears to be significantly less electropositive than the free guanidinium moiety present in GMQ and GBZ (Fig. 1B) and it is therefore likely to be less able to serve as a H-bond donor during its recognition at the ASIC3, which could underlie its differential response.

Given the overall 3D and electrostatic similarity of GBZ with GMQ (Fig.1), it was perhaps unsurprising to observe that GBZ followed a similar mode of action to GMQ, being capable of activating rASIC3 at pH 7.4 (Fig. 2A and 3A) and inducing a non-desensitising inward current but showing a higher potency but less efficacy than GMQ (Fig. 4F). Based on our findings from the blind docking experiments against the rASIC3-closed homology model (Fig. 3D and 3E), this difference could be explained by the lack of interaction with the core residue E79 and the adjacent residues V425 and L273. Such interactions are observed for GMQ in our docking experiment (Fig. 3E) and have been previously shown to be important in the hydrophobic interaction of the 4-methylquinazoline moiety of GMQ [61]. This is plausible because, unlike GMQ, GBZ lacks a second aromatic moiety in an appropriate position to allow such hydrophobic contacts to be established. Moreover, the sensitisation of the pH 6-induced rASIC3 activation observed by GBZ (Fig. 2D-G), but not by GMQ, could result from different residue interactions (Q271 and G269) of both molecules with rASIC3.

GBZ is an orally active α_2_-adrenoceptor agonist that has been used for many years as an antihypertensive drug [71]. However, GBZ also binds to the regulatory subunit of protein phosphatase 1, PPP1R15A, disrupting the stress-induced dephosphorylation of the α subunit of the translation initiation factor 2 (eIF2α), which protects against the detrimental accumulation of misfolded proteins in the endoplasmic reticulum (ER) [72] and has been proven effective in animal models that mimic misfolding protein diseases such as multiple sclerosis [73]. A recent study has identified a mono-chlorinated version of GBZ, sephin1, which retains the GBZ PPP1R15A inhibition activity, but without adrenoceptor activity [62] and it has also been shown to be effective in animal models of amyotrophic lateral sclerosis [62], Charcot-Marie-Tooth 1B (CMT1B) neuropathy [62] and multiple sclerosis [46]. Given the efficacy of sephin1 to ameliorate the pathogenesis of misfolding protein diseases, InFlectis Bioscience has begun the Phase I clinical trials to evaluate the safety of sephin1 (IFB-088) with the aim of evaluating its effect on the treatment of Charcot-Marie-Tooth 1B (CMT1B) disorder.

As a result of the similar pharmacological profile of GBZ to GMQ and with the intention of identifying a more selective rASIC3 modulator without adrenoceptor activity, we tested the effect of sephin1 on acid-induced rASIC3 activation. We observed that sephin1 shared a similar pharmacological profile with GBZ and GMQ, sensitising the response (sustained component) of rASIC3 to low pH (pH 6) (Fig. 4C and 4D) and mild acidosis (pH 7) (Fig. 5A-D), and activating rASIC3 at pH 7.4 (Fig.4E). These results were consistent with our expectations given that sephin1 appeared to be most similar to GMQ in terms of structural and chemical properties, exhibiting the highest TC score (Fig. 4A), and the highest molecular field points score (Fig. 4B). Moreover, increasing concentrations of sephin1 induced increasing potentiation of the transient and sustained phases of the pH 7-induced rASIC3 activation (Fig. 5A). In our blind docking experiments, carried out multiple times against three different ASIC3 models in open, closed and desensitized states, sephin1 poses tend to reproducibly cluster to the same site i.e. the nonproton ligand sensor domain where the GMQ and GBZ poses also cluster (Fig.7). Close inspection of the corresponding ligand interaction diagrams of sephin1 poses indicate that similar to GMQ and GBZ, the negatively charged residue E423 is likely to interact with the explicit guanidinium group of sephin1 in all three ASIC3 functional states (Fig. 6B, 6D and 6F). However, compared to GBZ, sephin1’s lack of a chloride atom in the aromatic ring seems to allow it to make hydrophobic interactions with surrounding residues like A378 (ASIC3-desensitised model, Fig. 6F), Q269 (ASIC3-open model, Fig. 6D) and L77 (rASIC3-closed model, Fig. 6B) from the adjacent subunit, but not with V425 as observed for GMQ (Fig.3E, 3G and 3I). Overall, these interactions likely underlie differential recognition of sephin1 at the nonproton ligand sensor domain of ASIC3, compared to that of GMQ and GBZ. GMQ is able to activate human ASIC3 (hASIC3) at neutral pH [74]. Given the structural and chemical similarities between GMQ, GBZ and sephin1, the close similarity between the nonproton ligand sensor domain in rASIC3 and hASIC3 (core residues E79 and E423, and adjacent residues, L77, Q269, L273, A378 and V425 are identical in both sequences [61]) and the predicted similar binding modes of all three molecules, it is likely that GBZ and sephin1 will also be able to modulate human ASIC3.

**Fig. 7.**
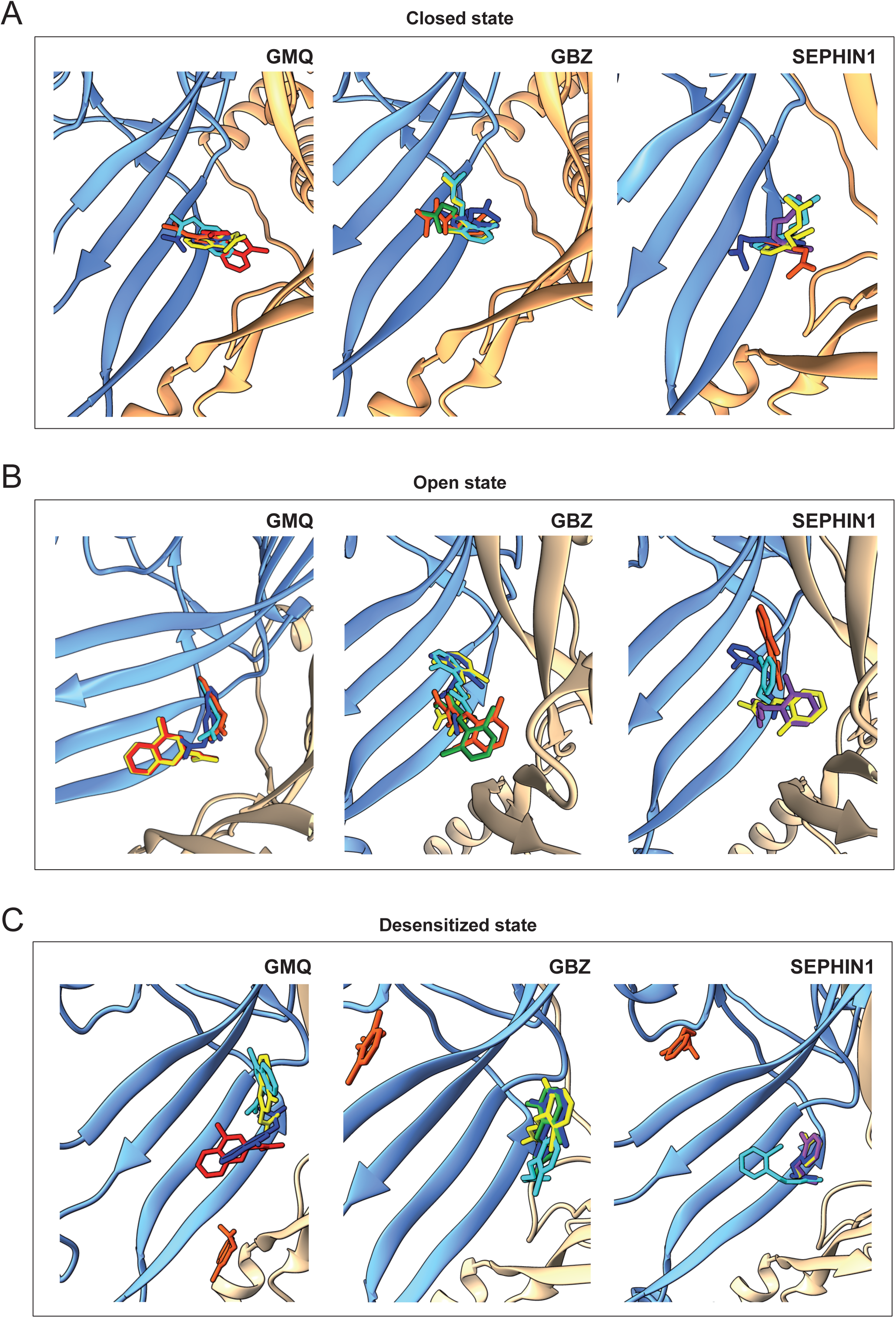
Docking of GMQ, GBZ and sephin1 to the rASIC3-closed, rASIC3-open and rASIC3-desensitized homology models. (A, C and E) Typical top 5 docked poses of selected drugs obtained through AutoDock 4.2.6-based blind docking against rASIC3 models that are likely to represent various states of the channel: rASIC3-closed (A), rASIC3-open (B) and rASIC3-desensitized (C). For clarity, only two subunits of the channel are shown in cartoon representations (subunit A in golden colour and subunit C in pale blue colour). For all panels, the top-ranked pose of GMQ, GBZ and Sephin1 are shown as red, green and purple, respectively. The second, third, fourth and fifth poses for each drug are shown in yellow, cyan, blue and orange red colour. Images were prepared using UCSF Chimera version 1.14.

Sephin1 is able to selectively disrupt the PPP1R15A-PP1c complex at 50 µM in cells *in vitro*, and after oral administration (1 or 10mg/kg) sephin1 accumulates in the nervous system reaching concentrations of up to 1 µM in the brain and sciatic nerve [62]. Moreover, a continuous 2-week treatment with sephin1 at 100 nM (once every 2 days) has shown efficacy in rescuing myelination of the dorsal root ganglia in a mouse model that mimics Charcot-Marie-Tooth 1B (CMT1B) in humans. Given the high functional expression of rASIC3 in DRG neurones and its involvement in pain derived from inflammatory diseases, together with the action of sephin1 observed on the channel, it is possible that sephin1 treatment could induce and/or exacerbate pain due to the activation of ASIC3 in DRG neurones, however our results show that both activation of rASIC3 at physiological pH (pH 7.4) and potentiation of the response to mild acidosis (pH 6) require higher concentrations of sephin1 (Fig. 4F and 5A) than those observed to be beneficial in treating misfolding protein diseases in mice. However, the possible additive effects of ASIC3 activated by sephin1 and other endogenous molecules found in the inflammatory soup such as arachidonic acid, which also potentiates the acid-induced response of ASIC3 [25] cannot be excluded. Moreover, ASICs function as trimers, displaying different pharmacological profiles depending upon subunit composition. In the present study, we have not evaluated the effect of sephin1 on heteromeric ASIC3-containing channels or other homomeric ASIC channels, and therefore, we cannot fully hypothesise what the potential effects of sephin1 *in vivo* might be. For instance, GMQ modulates ASIC1a and ASIC1b by shifting their pH dependence of activation to more acidic values [75], precisely the opposite effect seen for ASIC3, an effect attributed to structural differences in the extracellular domain of ASIC1a/b and ASIC3 [69]. Given the pharmacological and structural resemblance of sephin1 to GMQ, it is possible that sephin1 induces similar modulatory effects on other ASIC subunits. Nevertheless, we believe that the evaluation of pain thresholds in mice and humans used to study the effect of sephin1 in misfolding protein disease should be considered in future studies.

In summary, we have identified new rASIC3 modulators using a ligand-based *in silico* approach, namely GBZ and sephin1, and evaluated their effect on rASIC3 function using electrophysiology. Here we provide proof of principle, using a size-restricted chemical library (i.e. FDA-approved drug library), but believe this approach can be exploited in the future to screen much larger chemical space, enabling the identification of novel chemical scaffolds that act as ASIC modulators.

## Acknowledgements

The authors declare no competing financial interests. This work was supported by Versus Arthritis Research Grants (RG20930 and RG21973; GC and EStJS), BBSRC grant (BB/R006210/1; JRFH and EStJS), BBSRC-funded studentships (LAP and JCG, BB/M011194/1) and Gates Cambridge Trust (SC). TR gratefully acknowledges OpenEye Scientific Software and Cresset for granting academic licenses for some software used in the present study.

## Author Contributions

GC designed the research, conducted the experiments, acquired and analysed the data and wrote the manuscript. LAP, JCG, SC, EA and JRFH acquired and analysed the data. TR and EStJS designed the research and wrote the manuscript. All authors approved the final version of the manuscript.

## Declaration of Conflicting Interests

The authors have no conflicting interests to declare.

## Notes

#### Summary of Updates

New data to expand understanding of concentration-response effects and binding to different ion channel states, see figures 2, 4, 6 and 7.

## References

[1] L.A. Pattison, G. Callejo, E. St John Smith, Evolution of acid nociception: ion channels and receptors for detecting acid, Philos. Trans. R. Soc. B. 374 (2019). doi:10.1098/rstb.2019.0291.

[2] S. Kellenberger, L. Schild, International Union of Basic and Clinical Pharmacology. XCI. Structure, Function, and Pharmacology of Acid-Sensing Ion Channels and the Epithelial Na+ Channel., Pharmacol. Rev. 67 (2015) 1–35. doi:10.1124/pr.114.009225.

[3] R. Waldmann, G. Champigny, F. Bassilana, C. Heurteaux, M. Lazdunski, A proton-gated cation channel involved in acid-sensing, Nature. 386 (1997) 173–177.

[4] S. Gründer, M. Pusch, Biophysical properties of acid-sensing ion channels (ASICs), Neuropharmacology. 94 (2015) 9–18. doi:10.1016/j.neuropharm.2014.12.016.

[5] E. Deval, E. Lingueglia, Acid-Sensing Ion Channels and nociception in the peripheral and central nervous systems, Neuropharmacology. 94 (2015) 49–57. doi:10.1016/j.neuropharm.2015.02.009.

[6] S. Gründer, M. Pusch, Neuropharmacology Biophysical properties of acid-sensing ion channels (ASICs), Neuropharmacology. (2015) 1–10. doi:10.1016/j.neuropharm.2014.12.016.

[7] M. Hesselager, pH Dependency and Desensitization Kinetics of Heterologously Expressed Combinations of Acid-sensing Ion Channel Subunits, J. Biol. Chem. 279 (2004) 11006–11015.

[8] J. Jasti, H. Furukawa, E.B. Gonzales, E. Gouaux, Structure of acid-sensing ion channel 1 at 1.9 A resolution and low pH., Nature. 449 (2007) 316–23. doi:10.1038/nature06163.

[9] L.-N. Schuhmacher, G. Callejo, S. Srivats, E.S.J. Smith, Naked mole-rat acid-sensing ion channel 3 forms nonfunctional homomers, but functional heteromers, J. Biol. Chem. 293 (2018) 1756–1766. doi:10.1074/jbc.M117.807859.

[10] L.-N. Schuhmacher, E.S.J. Smith, Expression of acid-sensing ion channels and selection of reference genes in mouse and naked mole rat, Mol. Brain. 9 (2016) 97. doi:10.1186/s13041-016-0279-2.

[11] G. Callejo, A. Castellanos, M. Castany, A. Gual, C. Luna, M.C. Acosta, J. Gallar, J.P. Giblin, X. Gasull, Acid-sensing ion channels detect moderate acidifications to induce ocular pain, Pain. 156 (2015) 483–495. doi:10.1097/01.j.pain.0000460335.49525.17.

[12] E. Deval, J. Noël, N. Lay, A. Alloui, S. Diochot, V. Friend, M. Jodar, M. Lazdunski, E. Lingueglia, ASIC3, a sensor of acidic and primary inflammatory pain, EMBO J. 27 (2008) 3047–3055.

[13] S. Diochot, A. Baron, M. Salinas, D. Douguet, S. Scarzello, A.-S. Dabert-Gay, D. Debayle, V. Friend, A. Alloui, M. Lazdunski, E. Lingueglia, Black mamba venom peptides target acid-sensing ion channels to abolish pain, Nature. (2012) 1–24.

[14] E. Deval, J. Noël, X. Gasull, A. Delaunay, A. Alloui, V. Friend, A. Eschalier, M. Lazdunski, E. Lingueglia, Acid-sensing ion channels in postoperative pain, J. Neurosci. 31 (2011) 6059–6066.

[15] S. Kang, J.H. Jang, M.P. Price, M. Gautam, C.J. Benson, H. Gong, M.J. Welsh, T.J. Brennan, Simultaneous Disruption of Mouse ASIC1a, ASIC2 and ASIC3 Genes Enhances Cutaneous Mechanosensitivity, PLoS One. 7 (2012) e35225.

[16] A.J. Page, S.M. Brierley, C.M. Martin, C. Martinez-Salgado, J. a. Wemmie, T.J. Brennan, E. Symonds, T. Omari, G.R. Lewin, M.J. Welsh, L.A. Blackshaw, The ion channel ASIC1 contributes to visceral but not cutaneous mechanoreceptor function, Gastroenterology. 127 (2004) 1739–1747. doi:10.1053/j.gastro.2004.08.061.

[17] M.P. Price, G.R. Lewin, S.L. McIlwrath, C. Cheng, J. Xie, P. a Heppenstall, C.L. Stucky, a G. Mannsfeldt, T.J. Brennan, H. a Drummond, J. Qiao, C.J. Benson, D.E. Tarr, R.F. Hrstka, B. Yang, R. a Williamson, M.J. Welsh, The mammalian sodium channel BNC1 is required for normal touch sensation., Nature. 407 (2000) 1007–11. doi:10.1038/35039512.

[18] M.P. Price, S.L. McIlwrath, J. Xie, C. Cheng, J. Qiao, D.E. Tarr, K. a Sluka, T.J. Brennan, G.R. Lewin, M.J. Welsh, The DRASIC cation channel contributes to the detection of cutaneous touch and acid stimuli in mice., Neuron. 32 (2001) 1071–83.

[19] M. Mazzuca, C. Heurteaux, A. Alloui, S. Diochot, A. Baron, N. Voilley, N. Blondeau, P. Escoubas, A. Gélot, A. Cupo, A. Zimmer, A.M. Zimmer, A. Eschalier, M. Lazdunski, A tarantula peptide against pain via ASIC1a channels and opioid mechanisms, Nat. Neurosci. 10 (2007) 943–945.

[20] P. Zhang, Single Channel Properties of Rat Acid-sensitive Ion Channel-1alpha, -2a, and -3 Expressed in Xenopus Oocytes, J. Gen. Physiol. 120 (2002) 553–566. doi:10.1085/jgp.20028574.

[21] L.-N. Schuhmacher, S. Srivats, E.S.J. Smith, Structural Domains Underlying the Activation of Acid-Sensing Ion Channel 2a, Mol. Pharmacol. 87 (2015) 561–571. doi:10.1124/mol.114.096909.

[22] A. Baron, E. Lingueglia, Pharmacology of acid-sensing ion channels - Physiological and therapeutical perspectives., Neuropharmacology. (2015) 1–17. doi:10.1016/j.neuropharm.2015.01.005.

[23] R. Waldmann, F. Bassilana, J. de Weille, G. Champigny, C. Heurteaux, M. Lazdunski, Molecular Cloning of a Non-inactivating Proton-gated Na+ Channel Specific for Sensory Neurons, J. Biol. Chem. 272 (1997) 20975–20978. doi:10.1074/jbc.272.34.20975.

[24] J. Yagi, H.N. Wenk, L. a Naves, E.W. McCleskey, Sustained currents through ASIC3 ion channels at the modest pH changes that occur during myocardial ischemia., Circ. Res. 99 (2006) 501–9. doi:10.1161/01.RES.0000238388.79295.4c.

[25] E.S. Smith, H. Cadiou, P. a. McNaughton, Arachidonic acid potentiates acid-sensing ion channels in rat sensory neurons by a direct action, Neuroscience. 145 (2007) 686–698. doi:10.1016/j.neuroscience.2006.12.024.

[26] X. Wang, W.-G. Li, Y. Yu, X. Xiao, J. Cheng, W.-Z. Zeng, Z. Peng, M. Xi Zhu, T.-L. Xu, Serotonin facilitates peripheral pain sensitivity in a manner that depends on the nonproton ligand sensing domain of ASIC3 channel., J. Neurosci. 33 (2013) 4265–79. doi:10.1523/JNEUROSCI.3376-12.2013.

[27] T.W. Sherwood, C.C. Askwith, Dynorphin opioid peptides enhance acid-sensing ion channel 1a activity and acidosis-induced neuronal death., J. Neurosci. 29 (2009) 14371–80. doi:10.1523/JNEUROSCI.2186-09.2009.

[28] D.C. Immke, E.W. McCleskey, Lactate enhances the acid-sensing Na+ channel on ischemia-sensing neurons., Nat. Neurosci. 4 (2001) 869–70. doi:10.1038/nn0901-869.

[29] Y. Yu, Z. Chen, W.-G. Li, H. Cao, E.-G. Feng, F. Yu, H. Liu, H. Jiang, T.-L. Xu, A nonproton ligand sensor in the acid-sensing ion channel., Neuron. 68 (2010) 61–72. doi:10.1016/j.neuron.2010.09.001.

[30] W.-G. Li, Y. Yu, Z.-D. Zhang, H. Cao, T.-L. Xu, ASIC3 channels integrate agmatine and multiple inflammatory signals through the nonproton ligand sensing domain., Mol. Pain. 6 (2010) 88. doi:10.1186/1744-8069-6-88.

[31] S. Marra, R. Ferru-clément, V. Breuil, A. Delaunay, M. Christin, V. Friend, S. Sebille, C. Cognard, T. Ferreira, C. Roux, L. Euller-ziegler, J. Noel, E. Lingueglia, E. Deval, Non-acidic activation of pain-related Acid-Sensing Ion Channel 3 by lipids, EMBO J. 35 (2016) 1–15. doi:10.15252/embj.201592335.

[32] B. Cristofori-Armstrong, L.D. Rash, Acid-sensing ion channel (ASIC) structure and function: Insights from spider, snake and sea anemone venoms, Neuropharmacology. 127 (2017) 173–184. doi:10.1016/j.neuropharm.2017.04.042.

[33] S. Diochot, A. Baron, L.D. Rash, E. Deval, P. Escoubas, S. Scarzello, M. Salinas, M. Lazdunski, A new sea anemone peptide, APETx2, inhibits ASIC3, a major acid-sensitive channel in sensory neurons., EMBO J. 23 (2004) 1516–25. doi:10.1038/sj.emboj.7600177.

[34] W.S. Hsieh, C.C. Kung, S.L. Huang, S.C. Lin, W.H. Sun, TDAG8, TRPV1, and ASIC3 involved in establishing hyperalgesic priming in experimental rheumatoid arthritis, Sci. Rep. 7 (2017) 1–14. doi:10.1038/s41598-017-09200-6.

[35] M. Ikeuchi, S.J. Kolker, L.A. Burnes, R.Y. Walder, K.A. Sluka, Role of ASIC3 in the primary and secondary hyperalgesia produced by joint inflammation in mice., Pain. 137 (2008) 662–9. doi:10.1016/j.pain.2008.01.020.

[36] M. Ikeuchi, S.J. Kolker, K.A. Sluka, Acid-sensing ion channel 3 expression in mouse knee joint afferents and effects of carrageenan-induced arthritis., J. Pain. 10 (2009) 336–42. doi:10.1016/j.jpain.2008.10.010.

[37] M. Izumi, M. Ikeuchi, Q. Ji, T. Tani, Local ASIC3 modulates pain and disease progression in a rat model of osteoarthritis., J. Biomed. Sci. 19 (2012) 77. doi:10.1186/1423-0127-19-77.

[38] R.C.W. Jones, E. Otsuka, E. Wagstrom, C.S. Jensen, M.P. Price, G.F. Gebhart, Short-term sensitization of colon mechanoreceptors is associated with long-term hypersensitivity to colon distention in the mouse., Gastroenterology. 133 (2007) 184–94. doi:10.1053/j.gastro.2007.04.042.

[39] C. Reimers, C.-H. Lee, H. Kalbacher, Y. Tian, C.-H. Hung, A. Schmidt, L. Prokop, S. Kauferstein, D. Mebs, C.-C. Chen, S. Grunder, Identification of a cono-{RFamide} from the venom of {Conus} textile that targets {ASIC}3 and enhances muscle pain., Proc. Natl. Acad. Sci. U. S. A. 114 (2017) E3507–E3515. doi:10.1073/pnas.1616232114.

[40] K.A. Sluka, R. Radhakrishnan, C.J. Benson, J.O. Eshcol, M.P. Price, K. Babinski, K.M. Audette, D.C. Yeomans, S.P. Wilson, ASIC3 in muscle mediates mechanical, but not heat, hyperalgesia associated with muscle inflammation, Pain. 129 (2007) 102–112.

[41] A.A. Staniland, S.B. McMahon, Mice lacking acid-sensing ion channels (ASIC) 1 or 2, but not ASIC3, show increased pain behaviour in the formalin test., Eur. J. Pain. 13 (2009) 554–63. doi:10.1016/j.ejpain.2008.07.001.

[42] K. a Sluka, L. a Rasmussen, M.M. Edgar, J.M. O’Donnell, R.Y. Walder, S.J. Kolker, D.L. Boyle, G.S. Firestein, Acid-sensing ion channel 3 deficiency increases inflammation but decreases pain behavior in murine arthritis., Arthritis Rheum. 65 (2013) 1194–202. doi:10.1002/art.37862.

[43] D.J. Scholz, M.A. Janich, U. Kollisch, R.F. Schulte, J.H. Ardenkjaer-Larsen, A. Frank, A. Haase, M. Schwaiger, M.I. Menzel, Quantified pH imaging with hyperpolarized (13) C-bicarbonate., Magn. Reson. Med. 73 (2015) 2274–2282. doi:10.1002/mrm.25357.

[44] A.J. Wright, Z.M.A. Husson, D.-E. Hu, G. Callejo, K.M. Brindle, E.S.J. Smith, Increased hyperpolarized [1-(13) C] lactate production in a model of joint inflammation is not accompanied by tissue acidosis as assessed using hyperpolarized (13) C-labelled bicarbonate., NMR Biomed. 31 (2018) e3892. doi:10.1002/nbm.3892.

[45] F.S.J. Tennant, R.A. Rawson, Guanabenz acetate: a new, long-acting alpha-two adrenergic agonist for opioid withdrawal., NIDA Res. Monogr. 49 (1984) 338–343.

[46] Y. Chen, J.R. Podojil, R.B. Kunjamma, J. Jones, M. Weiner, W. Lin, S.D. Miller, B. Popko, Sephin1, which prolongs the integrated stress response, is a promising therapeutic for multiple sclerosis., Brain. 142 (2019) 344–361. doi:10.1093/brain/awy322.

[47] N.M. O’Boyle, M. Banck, C.A. James, C. Morley, T. Vandermeersch, G.R. Hutchison, Open Babel: An open chemical toolbox., J. Cheminform. 3 (2011) 33. doi:10.1186/1758-2946-3-33.

[48] P.C.D. Hawkins, A.G. Skillman, A. Nicholls, Comparison of shape-matching and docking as virtual screening tools., J. Med. Chem. 50 (2007) 74–82. doi:10.1021/jm0603365.

[49] E. Pihan, L. Colliandre, J.-F. Guichou, D. Douguet, e-Drug3D: 3D structure collections dedicated to drug repurposing and fragment-based drug design., Bioinformatics. 28 (2012) 1540–1541. doi:10.1093/bioinformatics/bts186.

[50] P.C.D. Hawkins, A.G. Skillman, G.L. Warren, B.A. Ellingson, M.T. Stahl, Conformer generation with OMEGA: algorithm and validation using high quality structures from the Protein Databank and Cambridge Structural Database., J. Chem. Inf. Model. 50 (2010) 572–584. doi:10.1021/ci100031x.

[51] E. Naylor, A. Arredouani, S.R. Vasudevan, A.M. Lewis, R. Parkesh, A. Mizote, D. Rosen, J.M. Thomas, M. Izumi, A. Ganesan, A. Galione, G.C. Churchill, Identification of a chemical probe for NAADP by virtual screening., Nat. Chem. Biol. 5 (2009) 220–226. doi:10.1038/nchembio.150.

[52] T. Cheeseright, M. Mackey, S. Rose, A. Vinter, Molecular field extrema as descriptors of biological activity: definition and validation, J. Chem. Inf. Model. 46 (2006) 665–676. doi:10.1021/ci050357s.

[53] E.S.J. Smith, X. Zhang, H. Cadiou, P.A. McNaughton, Proton binding sites involved in the activation of acid-sensing ion channel ASIC2a, Neurosci. Lett. 426 (2007) 12–17.

[54] I. Dittert, J. Benedikt, L. Vyklický, K. Zimmermann, P.W. Reeh, V. Vlachová, Improved superfusion technique for rapid cooling or heating of cultured cells under patch-clamp conditions, J. Neurosci. Methods. 151 (2006) 178–185. doi:10.1016/j.jneumeth.2005.07.005.

[55] N. Yoder, C. Yoshioka, E. Gouaux, Gating mechanisms of acid-sensing ion channels., Nature. 555 (2018) 397–401. doi:10.1038/nature25782.

[56] I. Baconguis, C.J. Bohlen, A. Goehring, D. Julius, E. Gouaux, X-ray structure of acid-sensing ion channel 1-snake toxin complex reveals open state of a Na(+)-selective channel., Cell. 156 (2014) 717–29. doi:10.1016/j.cell.2014.01.011.

[57] T. Rahman, E.S.J. Smith, In silico assessment of interaction of sea anemone toxin APETx2 and acid sensing ion channel 3., Biochem. Biophys. Res. Commun. 450 (2014) 384–9. doi:10.1016/j.bbrc.2014.05.130.

[58] G.M. Morris, R. Huey, W. Lindstrom, M.F. Sanner, R.K. Belew, D.S. Goodsell, A.J. Olson, AutoDock4 and AutoDockTools4: Automated docking with selective receptor flexibility., J. Comput. Chem. 30 (2009) 2785–2791. doi:10.1002/jcc.21256.

[59] B. Iorga, D. Herlem, E. Barre, C. Guillou, Acetylcholine nicotinic receptors: finding the putative binding site of allosteric modulators using the “blind docking” approach., J. Mol. Model. 12 (2006) 366–372. doi:10.1007/s00894-005-0057-z.

[60] S. Vullo, S. Kellenberger, A molecular view of the function and pharmacology of acid-sensing ion channels., Pharmacol. Res. (2019) 104166. doi:10.1016/j.phrs.2019.02.005.

[61] Y. Yu, W.-G.G. Li, Z. Chen, H. Cao, H. Yang, H. Jiang, T.-L. Le Xu, Atomic level characterization of the nonproton ligand-sensing domain of ASIC3 channels, J. Biol. Chem. 286 (2011) 24996–25006. doi:10.1074/jbc.M111.239558.

[62] I. Das, A. Krzyzosiak, K. Schneider, L. Wrabetz, M. D’Antonio, N. Barry, A. Sigurdardottir, A. Bertolotti, Preventing proteostasis diseases by selective inhibition of a phosphatase regulatory subunit, Science (80-.). 348 (2015) 239–242. doi:10.1126/science.aaa4484.

[63] J.-H. Lin, C.-H. Hung, D.-S. Han, S.-T. Chen, C.-H. Lee, W.-Z. Sun, C.-C. Chen, Sensing acidosis: nociception or sngception?, J. Biomed. Sci. 25 (2018) 85. doi:10.1186/s12929-018-0486-5.

[64] C.-C. Chen, A. Zimmer, W.-H. Sun, J. Hall, M.J. Brownstein, A. Zimmer, A role for ASIC3 in the modulation of high-intensity pain stimuli., Proc. Natl. Acad. Sci. U. S. A. 99 (2002) 8992–7. doi:10.1073/pnas.122245999.

[65] W.-N. Chen, C.-C. Chen, Acid mediates a prolonged antinociception via substance P signaling in acid-induced chronic widespread pain., Mol. Pain. 10 (2014) 30. doi:10.1186/1744-8069-10-30.

[66] J. Karczewski, R.H. Spencer, V.M. Garsky, A. Liang, M.D. Leitl, M.J. Cato, S.P. Cook, S. Kane, M.O. Urban, Reversal of acid-induced and inflammatory pain by the selective ASIC3 inhibitor, APETx2., Br. J. Pharmacol. 161 (2010) 950–60. doi:10.1111/j.1476-5381.2010.00918.x.

[67] M.G. Blanchard, L.D. Rash, S. Kellenberger, Inhibition of voltage-gated Na(+) currents in sensory neurones by the sea anemone toxin APETx2., Br. J. Pharmacol. 165 (2012) 2167–77. doi:10.1111/j.1476-5381.2011.01674.x.

[68] O. Alijevic, H. Hammoud, A. Vaithia, V. Trendafilov, M. Bollenbach, M. Schmitt, F. Bihel, S. Kellenberger, Heteroarylguanidines as Allosteric Modulators of ASIC1a and ASIC3 Channels, ACS Chem. Neurosci. 9 (2018) 1357–1365. doi:10.1021/acschemneuro.7b00529.

[69] T. Besson, E. Lingueglia, M. Salinas, Pharmacological modulation of Acid-Sensing Ion Channels 1a and 3 by amiloride and 2-guanidine-4-methylquinazoline (GMQ), Neuropharmacology. 125 (2017) 429–440. doi:10.1016/j.neuropharm.2017.08.004.

[70] X. Xiao, M.X. Zhu, T. Le Xu, 2-Guanidine-4-methylquinazoline acts as a novel competitive antagonist of A type γ-aminobutyric acid receptors, Neuropharmacology. 75 (2013) 126–137. doi:10.1016/j.neuropharm.2013.07.018.

[71] B. Holmes, R.N. Brogden, R.C. Heel, T.M. Speight, G.S. Avery, Guanabenz. {A} review of its pharmacodynamic properties and therapeutic efficacy in hypertension., Drugs. 26 (1983) 212–229.

[72] P. Tsaytler, H.P. Harding, D. Ron, A. Bertolotti, Selective inhibition of a regulatory subunit of protein phosphatase 1 restores proteostasis, Science. 332 (2011) 91–94. doi:10.1126/science.1201396.

[73] S.W. Way, J.R. Podojil, B.L. Clayton, A. Zaremba, T.L. Collins, R.B. Kunjamma, A.P. Robinson, P. Brugarolas, R.H. Miller, S.D. Miller, B. Popko, Pharmaceutical integrated stress response enhancement protects oligodendrocytes and provides a potential multiple sclerosis therapeutic., Nat. Commun. 6 (2015) 6532. doi:10.1038/ncomms7532.

[74] D. Barth, M. Fronius, Shear force modulates the activity of acid-sensing ion channels at low pH or in the presence of non-proton ligands., Sci. Rep. 9 (2019) 6781. doi:10.1038/s41598-019-43097-7.

[75] O. Alijevic, S. Kellenberger, Subtype-specific modulation of acid-sensing ion channel (ASIC) function by 2-guanidine-4-methylquinazoline., J. Biol. Chem. 287 (2012) 36059–70. doi:10.1074/jbc.M112.360487.

